# MELD-Adapt: On-the-Fly Belief Updating in Integrative Molecular Dynamics

**DOI:** 10.1101/2024.05.21.595263

**Authors:** Bhumika Singh, Arup Mondal, Kari Gaalswyk, Justin L MacCallum, Alberto Perez

## Abstract

Integrative structural biology synergizes experimental data with computational methods to elucidate the structures and interactions within biomolecules, a task that becomes critical in the absence of high-resolution structural data. A critical step in integrating the data, is knowing the expected accuracy or belief in the dataset. We previously showed that the Modeling Employing Limited Data (MELD) approach succeeds at predicting structures and finding the best interpretation of the data when the initial belief is equal or slightly lower than the real value. But, this initial belief is not known for many data sources. Here we introduce MELD-Adapt, designed to dynamically evaluate and infer the reliability of input data, thereby enhancing the accuracy of structure predictions. We demonstrate the utility of this method across different systems, particularly emphasizing its capability to correct initial assumptions and identify the correct fraction of data to produce reliable structural models. The approach is tested with two benchmark sets: the folding of twelve proteins with coarse physical insights and the binding of peptides with varying affinities to the Extrateterminal (ET) domain. Our findings also show the importance of how data is distributed in the system (locally clustered or globally distributed) and the subtle balance between data, beliefs, and force fields.

## Introduction

Integrative structural biology aims to synergize experimental insights with computational methods grounded in physical or statistical principles. The goal is to unravel the intricate structures and interactions within biomolecules and their complexes. Particularly invaluable in the absence of high-resolution structural data,^1^ these modeling tools address the challenge posed by biomolecules existing in multiple metastable states. In this regime, the main challenges arise from the combination of sparse structural data confounded by averaging of the signal over the diverse states present in the ensemble. Thus leading to ambiguous and sparse data, where the ambiguity arises from data that is compatible only with some states. But, the number of states to model is often unknown. In addition to the sparsity, ambiguity in the data, and an unknown number of states to model, the data is susceptible to random and systematic errors.^2^ In some instances, sampling of conformations is done independently of the data and the latter is used to reweight the ensembles and choose the most relevant biological states. This strategy works best when efficient sampling of phase space is accomplished, where the data helps to overcome sampling convergence or deficiencies in the scoring or physical function (e.g., force field) used. A second type of integrative approaches, and the focus of this work, is the use the data simultaneously with a sampling strategy to generate conformational ensembles. In such a scenario, the traditional integrative modeling workflow has four stages: 1) gathering information extracted from experiments; 2) selecting a representation of the system to model and mapping information onto the model, often as restraints; 3) structural sampling; and 4) model validation.^3^

Typically, the correct interpretation of the data is self-consistent with a particular structure, whereas random subsets of data are inconsistent with a structural and physical/statistical model. Thus, how much data an ISB approach believes is critical in determining the structures. If too little of the data is trusted, it leads to many possible subsets of data that are correct, but each of which is not very informative, reducing guiding power. On the other hand, when too much data is trusted, it increases guiding power, but it becomes incompatible with structural and physical/statistical models, producing incorrect predictions. These issues become exacerbated by the system’s internal dynamics and ensemble-averaged resolution of experimental data, leading to different possible interpretations of data, each compatible with different biologically relevant states. Thus, setting the data belief becomes critical.

To provide guiding power, the data is often transformed into some set of restraints (forward model) that the system has to satisfy, and which will incur some energy penalties when not satisfied. Some integrative approaches introduce a parameter with the fraction of data to trust, where the goal is to distinguish the informative data from the noisy one – and the structures compatible with it.^4–8^ In this scenario, satisfying restraints leads to no additional penalty energy or driving forces acting on the system, while violating restraint information leads to restraint energies and forces acting on the system, driving it to satisfy more data. However, fixing how much data to trust at the onset is based on average knowledge about techniques and could be an incorrect fraction for the current system. Enforcing too much data leads to inconsistent information that can result in inaccurate predictions while enforcing too little data reduces the possibilities of the dataset to direct towards the correct structure quickly. In other approaches, the accuracy value is not added. Instead, the functional form of restraints contributes constant energy beyond a certain threshold (e.g., step restraint potentials), which shifts the whole energy up. While they provide the advantage of foregoing the need to predefine a fraction of the data to trust, they also have less guiding power as there are no driving forces to satisfy more restraints.^9^ As integrative approaches continue to evolve, a critical need is for approaches that learn what fraction of the data is correct, which data to implement, and what structures are compatible with it. It is possible that different substates of the system, represented by the data, satisfy a different percentage of the data. To address these challenges, we introduce a novel Bayesian inference approach combined with MELD (Modeling Employing Limited data), called MELD-Adapt, designed to dynamically learn and adjust the trustworthiness of data inputs, thereby enabling more accurate structure predictions.

MELD^7,8^ uses a molecular dynamics (MD) sampling strategy at its core. MD approaches sample ensembles of conformations, which naturally leads to the identification of different states by analyzing the ensembles using the principles of statistical mechanics.^10^ However, MD is limited in its own way by the accuracy of the force fields and the ability to sample the diverse range of states that might be accessible to the system.^11^ For example, a conventional MD simulation in the microsecond timescale of a typical protein starting from an unfolded state will rarely visit the native state.^12,13^ Leveraging data in MD through MELD focuses sampling on regions of phase space compatible with data while simultaneously identifying the best interpretation of the data compatible with the physical model. Solving these two problems together reduces the computational time needed to sample biologically relevant states, and can in some cases compensate for deficiencies in the force field or modeled system (e.g., when having a partial model such as solving the structure of a protein that has a ligand in the experiment but not in the simulation)^14^. The difficulty lies in knowing the amount of data from the available pool that should be used. It is especially important when combining sources of data that might mitigate individual limitations in each set.^15^ When dealing with new sources of data, we often need to run with different protocols to identify a robust approach where not too much or too little information is used. But this rapidly increases computational expense. And, in many cases is not transferable among systems where the same type of data is used.

In this work, we introduce a Bayesian inference approach overlaid with the MELD method to address the challenges of unknown uncertainties in experimental datasets. This development allows users to provide a prior estimate of the amount of data to trust for a specific data type, while MELD identifies the actual amount of data it should trust. We demonstrate the effectiveness of this approach across different systems for protein and protein-peptide structure prediction, showcasing its ability to predict accurate structures even when initial assumptions are incorrect. First, we show that when the initial data accuracy is correct (e.g., has already been optimized), MELD-Adapt is able to predict accurate structures in agreement with MELD. Then we show that even when the initial assumptions are incorrect, MELD-Adapt is able to identify the correct fraction of data and the structures compatible with it, whereas traditional MELD makes incorrect predictions.

**Figure 1:**
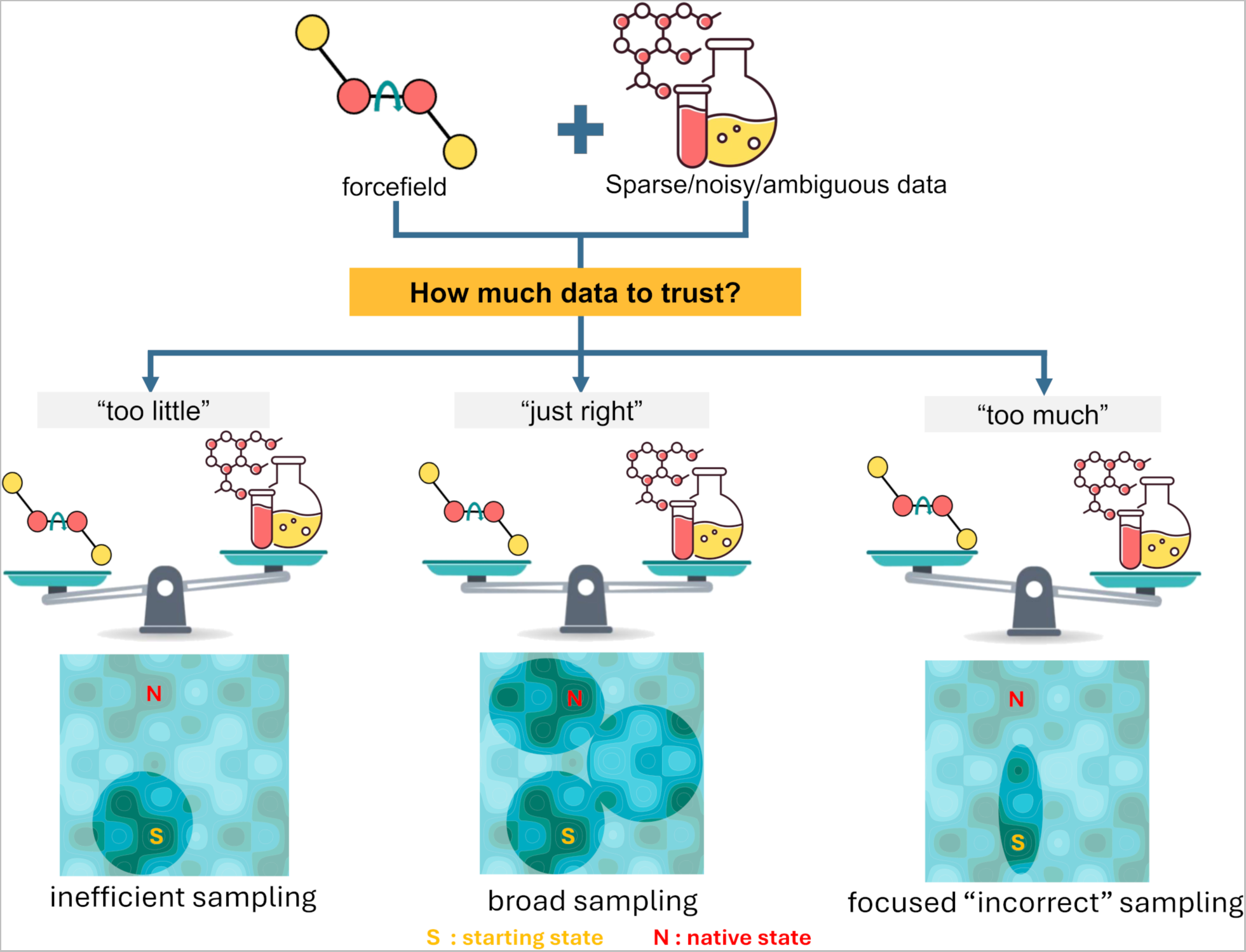
Balancing the quantity of semi-reliable data in guiding simulations: Insufficient data leads to inefficient sampling (left), while an excess of inaccurate data can result in highly focused yet incorrect predictions (right). Finding the optimal balance between data and physics-based models ensures broad and accurate sampling (middle). The letters S and N represent the starting and native states, respectively

## Computational Methods

### The Modeling Employing Limited Data (MELD) approach

The MELD methodology has been previously described^7,8^ and thus only a quick summary is provided here. The MELD philosophy combines information that might be ambiguous and noisy with molecular simulations through Bayesian inference. Ambiguity refers to a source of data that might have multiple implementations where only one is correct in the native structure of the biomolecules (e.g., atoms A and B are within 5 Å of each other or A and C are within 5 Å). By noisy data, we mean that for some data none of the possible interpretations are found in the native structure.

MELD does not use all data simultaneously to accommodate for noise and ambiguity. Rather, we provide an accuracy value for each dataset (a collection) that determines how many data points should be enforced, but not which data points. Which data is enforced dynamically changes throughout the simulation. A simulation can have multiple collections originating from different experiments or data sources. During the simulation, all of the data points in a collection are evaluated and ranked by the restraint penalty they introduce to the simulation. Then, the data points with the lowest energy are used until the next simulation step – driving the dynamics together with the force field. How many data points are used is fixed and determined by the accuracy value and number of data points in the collection. These choices provide a deterministic way for selecting which data points to enforce given a certain sampled structure.

MELD sampling at low temperatures, where restraints are strongly enforced, rarely changes which restraints are active, as this would require crossing over large energy barriers. To facilitate sampling of different subsets of data that guide to different regions of phase space, we use a Hamiltonian and Temperature Replica Exchange ladder.^16^ At the highest replicas, temperature is high, and force constants for restraints vanish, allowing the efficient sampling of phase space and enforcing different subsets of data. As those structures are exchanged to lower replicas, the subset of restraints with the lowest restraint energy becomes active, guiding sampling towards regions of phase compatible with the active subset of data.

In the end, the posterior distribution agrees with both the force field and the best interpretation of the data. Framed as a Bayesian inference approach (see Equation. 1), where x represents a particular conformation at a timestep given by an atomistic force field and D represents corresponding NMR data. The prior (*p*(*x*)) originates from the Boltzmann distribution given a force field and the data likelihood (*p*(*D*|*x*)) is given by the energy penalty of using some subset of data given the sampled structure. Finally, *p*(*x*|*D*) is the posterior distribution from which we sample. It represents the probability of sampling a specific conformation given the fraction of the data enforced in the simulation.

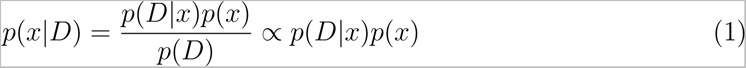

The critical step here is to determine the accuracy value for each collection. Users need to develop a deep understanding of the technique and system before selecting useful values for this – often having to compare agreements between different protocols using different accuracy values to ensure consistency and robustness of the approach. In the next section, we introduce a new implementation that allows the simulation to learn the accuracy value through the simulation using Bayesian inference.

### Inferring the accuracy value for a collection: MELD-Adapt

A fixed (static) accuracy value forces the decision upfront. If the user chooses a value that is too high, the guarantees MELD has about the relevance of the most populated state in the posterior ensemble can be incorrect. On the other hand, choosing a number much lower than the dataset accuracy implies losing directive power in the simulations. Ideally, the accuracy can dynamically change during the simulation. This implies that throughout the simulation, in addition to selecting the most appropriate subset of data to use, there is also a process to ascertain the optimal amount of data to be utilized. To address this limitation we have extended the Bayesian inference approach to consider the likelihood of enforcing the data (*D*) given a structure (*x*) and a number of active restraints (*y*). Thus, the posterior probability becomes:

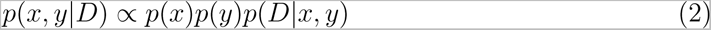

Where *p*(*x*) represents the prior probability over the structural variable x and is modeled using the Boltzmann distribution generated by the forcefield; *p*(*y*), reflects the prior probability over the number of active restraints *y*, which crucially determines the number of active restraints associated with each collection during the simulation; and *p*(*D*|*x*, *y*), which quantifies the likelihood of observing a subset of the given data D under a specific structural conformation *x* and a particular set of active parameters *y*. Our primary focus for structural determination typically centers on the marginal distribution *p*(*x*|*D*), which is obtained by integrating out the variable *y* to provide a more comprehensive understanding of the structural aspects of the system, all driven by the available data. Although *x* and *y* are dependent on each other (*p*(*D*|*x*, *y*) is evaluated as a log-likelihood involving both variables), their priors (*p*(*x*) and *p*(*y*)) are independent as one originates from the force field (*p*(*x*)) and the other from the forward-model used to implement the experimental data (*p*(*y*)). We next consider what constitutes a good prior for the data (*p*(*y*)).

Restraints are typically enforced as flat-bottom harmonic potentials, implying that their energy contribution is always greater or equal to zero. Thus, an on-the-fly increase in the number of restraints to satisfy would typically imply an increase in energy (unless a restraint is already satisfied, in which case it would not bring additional directive power). On the other hand, reducing the number of restraints will typically lead to a reduction in the overall restraint energy. Consequently, a naive approach would have a strong tendency towards satisfying zero restraints at any point of the simulation and thus lose any benefit MELD brings.

We employ a dynamic approach to update the active fraction of restraints throughout the MD simulation by utilizing Monte Carlo (MC) trials. We initialize the simulation with a prior for each collection based on the expected accuracy and a maximum and minimum range. We perform a Monte Carlo trial at given simulation intervals (e.g., every 100th MD step). During each trial, the number of active restraints can change by up to ±5. If, after the trial, the system has a number of restraints between the maximum and minimum values, the change in the number of restraints is accepted based on the Metropolis criterion. In the Metropolis criterion decreasing the energy will always result in acceptance of the change. Since the restraint energy for each restraint is always greater or equal to zero (*E_rest_* ≥ 0), reducing the number of restraints would always be favored. In essence, this corresponds to having a uniform prior that does not promote or discourage a specific number of restraints and just relies on the restraint energy. As the restraint energy with a uniform prior would lead to satisfying few restraints and therefore losing guiding power, it becomes important to establish a prior that promotes enforcing a higher number of restraints.

We decided to use an exponential prior to reward the addition of restraints over the initial value (see Fig. 2). Effectively, this prior is introduced by adding a reward energy term for every restraint above an initial value, the *E_reward_* = −λ*k_B_T* (*N_i_* − *N_prior_*), where λ determines the magnitude of the reward per restraint, *N_i_* is the number of current proposed restraints and *N_prior_* the number of restraints we set as a prior for the simulation. Effectively, this *E_reward_* is the log-prior and requires two hyperparameters: λ (reward magnitude) and *N_prior_* (starting belief). When λ is set to zero, we return to the uniform distribution prior. When λ *>* 0, we have an exponential prior. When combining the prior with the restraint energy and force field energy, we obtain a distribution of sampled restraints (see Fig. 2). As λ increases, the method would enforce more and more data – overcoming force field preferences. Thus, it is important to find a good reward magnitude that will balance between the best use of the data and forcefield. Below we detail the type of data used in this study and how we identified the value for λ.

**Figure 2:**
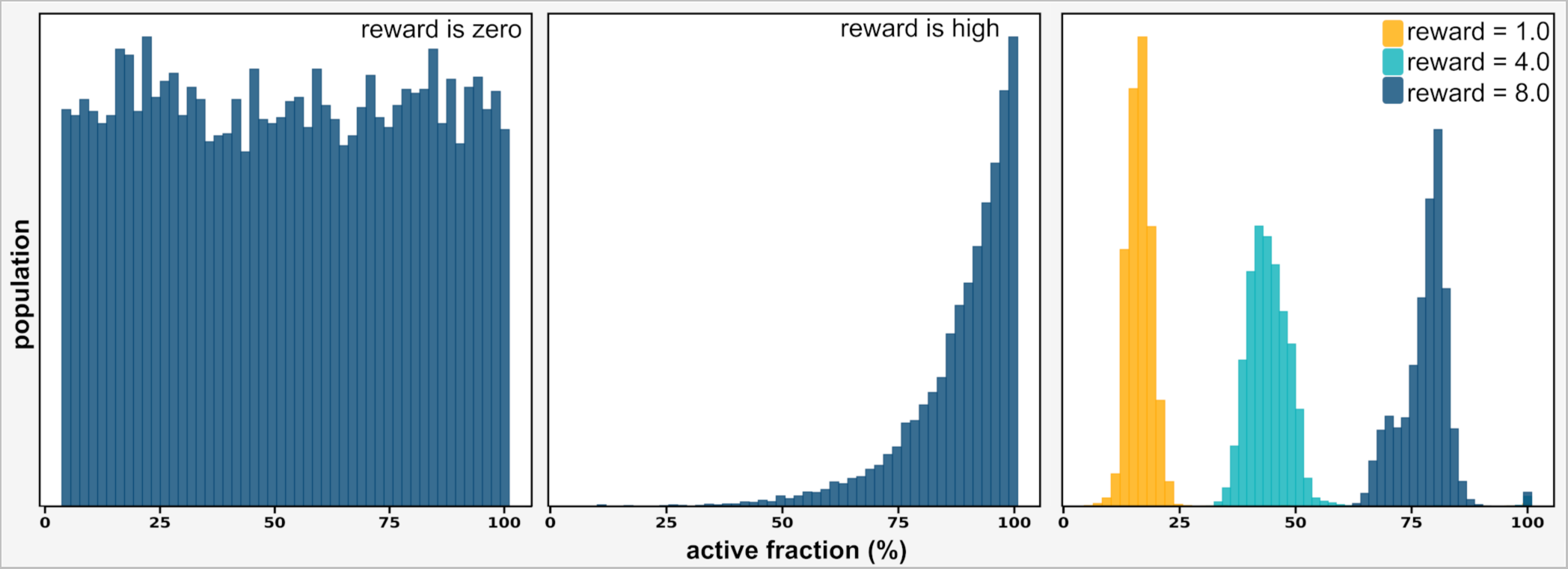
Effect of reward value on prior distribution. When the reward is zero, the prior is constant, and it uniformly samples all possible numbers of restraints in the absence of any forces(left). When the reward is non-zero, the prior grows exponentially, favoring a large number of active fractions in the absence of any forces (middle). In the presence of a force field, the active fractions find an upper and lower limit governed by the reward value and the force field (right).

### Benchmark datasets

We use two benchmark sets for which we have previous experience using MELD with fixed accuracy values. The first benchmark is the folding of twelve proteins starting from sequence using coarse physical insights (CPIs) as external information. The second benchmark is the binding of a series of peptides that fold upon binding the Extrateterminal (ET) domain of Bromo and Extrateterminal domain (BET) proteins using chemical shift perturbation (CSP) data for the ET domain measured in the presence/absence of the peptide. Each type of data (CPIs or CSPs) has its own strengths and weaknesses.

CPIs originate from general protein principles^8^ such as the presence of hydrophobic cores or pairing of beta strands based on secondary structure predictions using PSIPRED protein structure prediction server.^17^ This leads to many possible restraints across pairs of residues along the sequence. Since this data does not originate from an experiment, it is typically less self-consistent than that from experiments and provides less guiding power. The advantage is that this data is easy to derive starting from the protein sequence, but the amount of possible restraints and, therefore, the amount of noise increases with protein complexity (in terms of secondary structure) and size. Thus, the approach has only been successful for small proteins.

CSP datasets for peptide binding provide information about chemical shifts in the protein that change in the presence/absence of the peptide but provide no structural information about the peptide. Typically, we process this data to identify residues in the protein with the largest perturbation and hypothesize that this change could be due to interactions with the peptide. However, we do not know which residues in the peptide interact, thus generating all possible interaction combinations between peptide residues and protein residues above the threshold, leading to a noisy dataset. Some of the selected protein residues might not even be involved in interactions with the peptide and change their chemical environment due to allosteric changes, further increasing the noise level in the dataset.

The major difference between CSP and CPIs is that CPIs are distributed through the 3D space of the protein, thus leading to many incompatible sets of restraints. On the other hand, CSP data tends to be localized near the binding site; thus, small conformational changes in the protein-peptide interaction easily give rise to satisfying a higher number of restraints (e.g., an extended conformation wrapping around the active site) – challenging the biological relevance of the simulations.

### Simulation Details

#### Protein Folding Benchmark

We conducted MELD-Adapt simulations on a dataset comprising 12 proteins (listed in SI Table 1). All simulations were performed using the OpenMM^18^ suite of programs, starting from fully extended chains and a combination of the ff14SB^19^ and ff99SB^20^ atomic force fields for side chains and backbone, respectively. The simulations were executed within the GBNeck2 implicit solvent model,^21^ using hydrogen mass repartitioning and a 3.5 fs time step.^22^ To mitigate issues related to local energy minima, and to achieve efficient sampling, MELD uses Hamiltonian and Temperature Replica Exchange Molecular Dynamics (H, T-REMD).^16^ We used 30 replicas such that each replica operates at varying temperatures within the range of 300K to 550K while maintaining a strong force constant (250 KJ·mol·nm^−2^) for low-temperature replicas and gradually reducing it to zero for high-temperature replicas.

We performed simulations choosing different values for the two parameters that control how much data is enforced: the reward value and the prior belief. For the reward value, *E_reward_* = −λ*k_B_T*Δ, choosing higher rewards promotes the activation of a higher number of restraints. We experimented with different values of λ, specifically 0.25, 0.5, 1.0, 2.0, 4.0, and 8.0. Consequently, six different simulations were conducted for each test system, each utilizing different reward values. For these systems, we chose our initial belief as we had done in our prior work^8^– which had already been optimized to account for most proteins (each hydrophobic residue is on average in contact with 2.4 other hydrophobic residues) and 45% of residues predicted to be part of β-strands will be involved in N-H…O hydrogen bonding to another strand residue. We then selected three systems (3GB1, T0769, and T0773) for which we repeated the calculation with an initial belief that was either lower (1.2 hydrophobic contacts and 25% of strand residues directed towards strand pairing) or higher than our previous work (4.8 hydrophobic contacts and 65% of strand residues directed towards strand pairings).

#### Protein-peptide Binding Benchmark

We chose 5 different reward values to optimize: λ = [0, 0.25, 0.5, 0.75, 1.0]. Furthermore, for these systems, where intrinsically disordered peptides fold upon binding, the force field plays a crucial role. Thus, along with the reward value optimization, we tested there different force field and solvent model combinations: ff14SBside+gbNeck2,^19,21^ ff14SBside+ obc,^19,23^ ff12SB-cMAP+ obc^23,24^ – for each of them we ran the traditional MELD approach with fixed number of data belief or the current MELD-Adapt protocol. Our previous experience with this systems used standard MELD with the ff14SB+ gbNeck2 combination.^25^

Simulations start from an unstructured peptide far from the protein receptor. We use the ExtraTerminal domain of BRD3 with peptides TP, NSD3, CHD4, and BRG1 peptides and the ExtraTerminal domain of BRD4 for the LANA peptide based on the solved NMR structures deposited in the PDB (7JQ8, 7JYN, 6BGG, 6BGH, 2ND0).^26–28^ We used available CSP data from TP and NSD3^26^ and transferred the ET:TP data to the remaining systems, as previously done.^25^ We set our initial trust of the CSP data at 4% for both MELD and MELD-Adapt protocols. Each simulation ran for 1 µs with 30 replicas. Temperature and restraint strength were scaled non-linearly. Temperature ranged from 300K to 500K, and force constant ranged from 350 kJmol^−1^nm^−2^ in the lower replicas to 0 kJmol^−1^nm^−2^ at the highest replica. More detail regarding setting up H,T-REMD protocol can be found in our previous study.^25^

### Analysis

We analyze the ensembles produced by different protocols by their ability to predict native-like confirmations (e.g., in the highest population cluster), provide faster and more robust convergence to the native state (e.g., by looking at the behavior across all replicas), and by finding the ideal fraction of the data to trust. This leads to the following three types of analysis:

#### Clustering

We used hierarchical agglomerative clustering to group structurally similar configurations and determine their respective populations using AMBER’s *cpptraj* package.^29^ The clustering process was applied to the second half of the trajectory from the five lowest temperature replicas, utilizing a cutoff distance of 2 Å and considering only Cα carbons of residues with predicted secondary structure. For protein-peptide complexes, we used a cutoff of 1.5 Å for clustering, and unstructured peptide regions were removed from clustering. Our reported predictions include the centroid of the most populated cluster (top1) or the best centroid from the top 5 population clusters (top5) and their population and RMSD from native (interface RMSD or iRMSD^30^ for protein-peptide complexes). Additionally, we assessed the fraction of frames with structures that conformed to the same number of restraints as native structures.

#### Convergence checks

We conducted structure prediction convergence checks by analyzing the ensembles produced by the 30 replicas. First, we analyzed the RMSD distribution for the ensemble generated at each replica (the different Hamiltonian and Temperature). We anticipated observing RMSD distributions that would be wide at the highest replica and become narrower distributions (either native-like or misfolded) at lower replicas, in essence following a funnel-like behavior. We expect better protocols to yield a more funneled landscape toward native-like structures. Whereas bad protocols will converge quickly to non-native states.

Second, we checked convergence by comparing RMSD distributions against the native state for each individual walker as it performs a random walk in replica space. A well-converged simulation is characterized by consistent RMSD distributions among all walkers, with peaks occurring at the same values. Simulations that do not exchange properly (e.g., exchanging locally without achieving roundtrips in replica space) will present very different RMSD distributions. In a converged simulation with stable conformations, each walker should be able to visit native-like structures, with RMSD values below 4 Å. We quantified convergence using the Kullback-Leibler (KL) Divergence using the Scikit-learn package.^31^ A KL divergence value of 1.0 or higher implies a significant disparity with the other individual distributions, while a value of 0.0 indicates an exact match between the observed and the other individual distributions.

#### Comparison of MELD predicted accuracy with the amount of data satisfied by the experimental structure

MELD-Adapt simulations sample a distribution of accuracies in a replica-dependent manner. We expect the analysis of the distribution of enforced accuracy at the lowest replica index to overlap significantly with the number of restraints satisfied by the native structure. Thus, we analyzed the number of hydrophobic and strand pairing restraints (for protein folding) and CSP-derived distance restraints (for peptide binding) within the structure of each frame at the lowest temperature replica relative to the native structure using the MDTraj package. ^32^ The optimal reward value (λ) should yield a distribution that overlaps with the distribution obtained from satisfied restraints.

## Results and Discussion

MD-based approaches typically have a hard time disambiguating between sampling issues and force field issues. In this study, the addition of restraints can overpower force field preferences, and as we allow the number of restraints to change, so can the balance between force field and restraint energy contributions be skewed. This effect becomes especially pronounced in peptide-protein complexes, where the nature of the data and the marginal stability of the peptide, with the balance between intrinsically disordered proteins/peptides (IDPs) (unbound) and folded (bound), is more sensitive to changes in the number of enforced restraints. Thus, we will present and discuss the two benchmark sets separately and then provide a broader discussion.

### MELD-Adapt samples folded states across a wide range of protocols

For each system and simulation condition, we look at the ability of the approach to 1) sample the native state, 2) identify the native state as the highest population cluster, 3) identify the correct interpretation of the original data, and 4) agree between replica walkers (convergence). Figure 3 summarizes our results across different protocols for Protein G (PDB ID 3GB1). First, we look at the funneling power of different protocols, that is, the ability to narrow down sampling in low-temperature replicas to focus on the native state. Second, we look at the overlap in RMSD distributions along different replicas and calculate the KL-divergence to assess convergence – if only one replica finds the native state for a long time the system is less converged than if all replicas sample the native state for a short amount of time. Third, we look at how much data (how many active restraints) the system is imposing during the simulation, and how many restraints are actually satisfied. If more restraints are enforced than satisfied, that means that we are adding restraint energy to our potential, which could alter force field preferences. On the other hand, if we are satisfying more restraints than actually enforced, this shows that the cooperative nature of restraints leads to satisfying more restrains than the ones being enforced (e.g., a contact between residues *i* and *j* trivially results in a contact between *i* and *j* +1 even if there is no restraint guiding to it). Fourth, we look at the RMSD value between the centroid of the top population cluster and the native structure.

**Figure 3:**
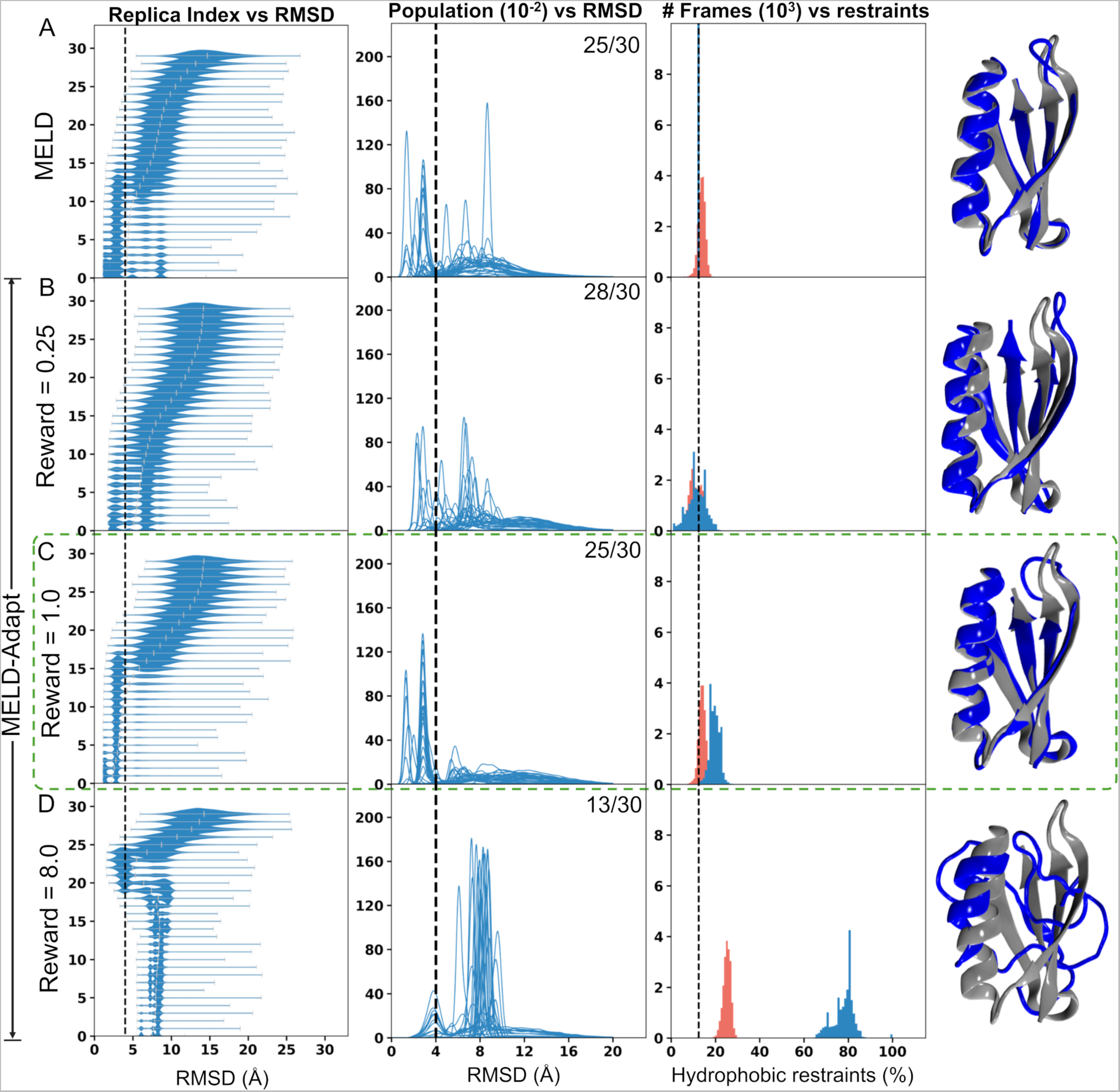
Comparative analysis of protocols with varying reward values for Protein G. Panel A uses a fixed belief with traditional MELD, while Panels B-D uses MELD-Adapt with different λ values. The first column shows the funneling power of each protocol; the second column - shows the overlap in RMSD distributions across 30 walkers, the values at the top-right indicate how many of the 30 replicas have an average pairwise KL divergence lower than 0.5. The third column - shows the percentage of active restraints during simulation (in blue) and the percentage of restraints satisfied (in red). The Fourth column shows the centroid of the top population cluster (in blue) compared to the native structure (PDB ID: 3GB1, in grey).

In general, when the data is not directive enough the funneling plots will exhibit broader distributions in the lowest temperature replica (see panels A, B in Fig. 3). They will also rapidly lose native-like populations as we go to higher replicas. On the other hand, when enough data is enforced this leads to narrow RMSD histograms in the lowest temperature replica (see panel C in Fig. 3). When a higher amount of data is enforced than the correct amount it can also lead to narrow distributions that are shifted with respect to the native state: the method guides to an incorrect region of phase space (see panel D in Fig. 3).

Even with the fixed accuracy in MELD, we observe a distribution along the number of restraints that are satisfied in the native state centered around the enforced value (see panel A in Fig 3). This is due to the way we introduce restraints using flat-bottom harmonic restraints, where there is a quadratic term extending one Å in each direction that contributes *k_B_T* at its maximum. Hence, thermal fluctuations can already account for small changes in satisfied restraints close to the fixed MELD value. Interestingly, with a reward value of one, we start seeing a separation between the number of restraints enforced (that is, contributing a restraint energy term to the potential, blue distribution) and those actually satisfied (red distribution, contributing an energy of 0 kcal/mol to the potential energy). Although protocols using a reward value between 0.25 and 1 kcal/mol per restraint activated above the initial belief systematically find the native state with high accuracy, the latter has a greater funneling potential and greater convergence of the results across replicas. As the difference between satisfied and enforced restraints becomes larger (λ = 4 and higher), we observe convergence towards incorrect regions of phase space.

Fig. S1-S10 shows the results across the 10 protein systems and set of conditions. Ubiquitin, the most complex protein in the data set and a slow folder is a failure across all protocols. With a reward value of 0.5 kcal/mol per restraint yielding the best results (top cluster centroid 6.5 Å from native). The failure in this system is mostly attributed to the small 3-10 helix which forms at the end of the simulation and is not correctly sampled in simulations. For the remaining nine systems, MELD, and MELD-Adapt with rewards between 0.25 and 1 all produce excellent results, where the native state is always found as a top 5 population cluster, and for the 0.25 and 0.5 rewards systems also in the top population cluster. However, looking at Fig. S11, it becomes evident that, in general, the higher reward value of 1 leads to higher populations for the top cluster. Surprisingly, there are no clear trends as to the effect of different protocols on first passage time (the first time we sample the native state) (see SI table 2). This is consistent with the two state-nature of many of these proteins, where forming a few critical contacts will directly lead to the native state. By activating different percentages of the data, the different protocols allow other possible states to be sampled but do not change the speed of each individual event.

### MELD-Adapt recovers from incorrect beliefs

While the previous results show the importance of choosing the correct reward value, they show similar results between MELD and MELD-Adapt. The CPI initial belief was already optimized when developing the CPI protocols. Thus, MELD-Adapt converges on the MELD solution. To showcase the power of MELD-Adapt over MELD, we prepared some simulations using an incorrect initial belief. We chose three systems (protein G, and CASP11^33^targets T0769 and T0773),^14^ where we set the initial belief to be substantially below or above the real accuracy of the data. For these three tests, we used our previously selected reward value of 1 kcal/mol. Fig. 4 and Fig. S13-14 summarize our results across these systems. The funneling plots show that, indeed, the MELD-Adapt method obtains similar results irrespective of the initial belief. This is a significant improvement over MELD, where enforcing too little data is not directive enough for protein G; and enforcing too much data directs protein G to an incorrect region of phase space. Surprisingly, for T0769 and T0773, we obtained good results irrespective of the initial belief and whether MELD or MELD-Adapt was used. Both T0769 and T0773 were protein designs from the Baker lab, which have been optimized to have very good funneling behavior and more stability than naturally occurring proteins. Thus, in this case, the balance between force field and restraint penalties favors the native state even when too much data is being enforced.

**Figure 4:**
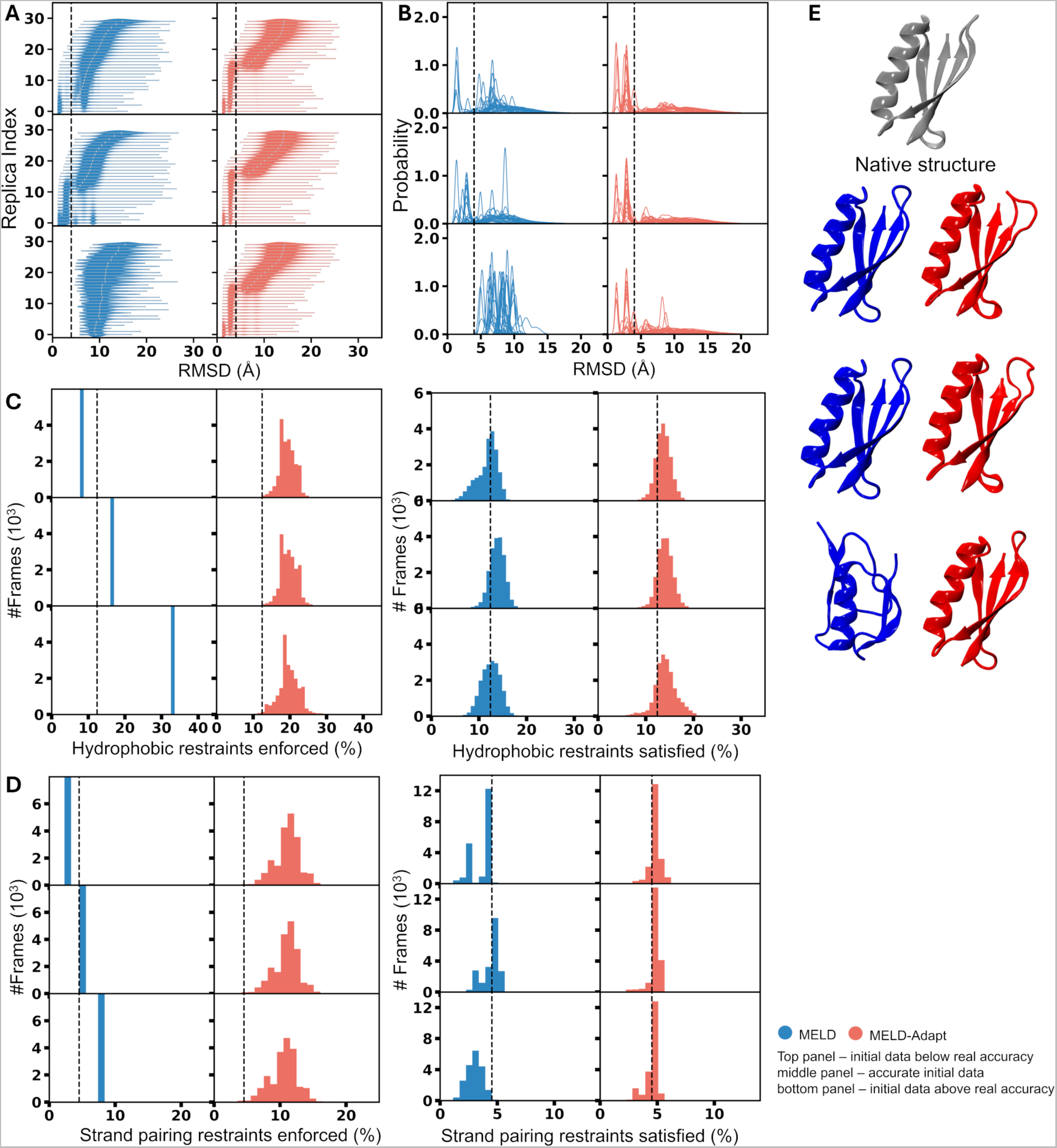
Folding Belief Analysis. Comparison of protein folding behavior between MELD (blue) and MELD-Adapt (red, with a reward value of 1.0 kcal/mol) under three scenarios: i) initial data below real accuracy, ii) accurate initial data, and iii) initial data above real accuracy. The analysis includes A) Funneling protocol, B) RMSD distribution, C) percentage of enforced and satisfied hydrophobic restraints, and D) percentage of enforced and satisfied strand pairing restraints, illustrating that MELD-Adapt dynamically adjusts the trust percentage and is independent from initial beliefs. E) Best predicted structures in each protocol compared to the native structure (silver).

### Peptide Binding

We chose a series of five peptide systems that bind the ET domain with different binding affinities ranging from nanomolar (90 nM) to sub-millimolar (635 *µ*M). We had previously shown that using CSP data alone we could predict the structure of the complex for the three strongest binders (TP, CHD4, and BRG1). For NSD3, the structure of the complex was within the top 5 clusters, and for LANA, it was within the top 10 clusters. One of the caveats in mapping the experimental data from CSP is that the sensitivity of the data is dependent on the binding affinity; hence, we had to decide on a different threshold for different systems. Furthermore, we used the CSP data from the TP and transferred it to use for BRG1, LANA, and CHD4. The threshold allows us to determine how many protein residues will likely be involved in the protein-peptide interface. Since there is no information for the peptide we have to consider that any residue in the peptide could be involved in binding – this leads to a combinatoric of possible contacts between protein and peptide. Since the peptides are of different lengths (see SI table S6), this also means that the number of total restraints is different for each system. In our previous work, we used a 4% accuracy after looking for self-consistency in predictions across simulations enforcing different belief values. Ideally, parameter sampling would allow us to recover the structure of the complex as the top prediction for all systems.

As in the case of folded proteins, we attempted different reward values, first starting from the ff14SBside force field and GBneck2 implicit solvent we had used in previous work (see second row in Figure 5). However, in this case, reward values above 0.50 kcal/mol rapidly lead to incorrect predictions where the peptide stretched engulfing the active site (See Figure S15). Upon reflection, all restraining contacts in the dataset for these systems are clustered together, and the peptides are at the threshold of stability for folding upon binding. Hence, even a small reward for satisfying more restraints can easily overcome force field preferences – enabled by the close proximity of all candidate restraints. As the reward value increases, we observe a rapid shift in enforced and satisfied restraints towards higher values that should be enforced (see Figure S16). With a reward of 0.25 kcal/mol, we noticed that for all peptides the predicted top structure had the peptide in the correct binding site – but the peptide conformations were incorrect for several peptides. TP, the strongest binder in the set, is correctly predicted irrespective of the protocol. LANA, the weakest binder in the set, was also predicted in the active site, but the interacting amino acids were shifted along the sequence with respect to the experimental structure. Finally, both CHD4 and BRG1 bound as strands, but in a flipped conformation (see third row in Figure 5).

**Figure 5:**
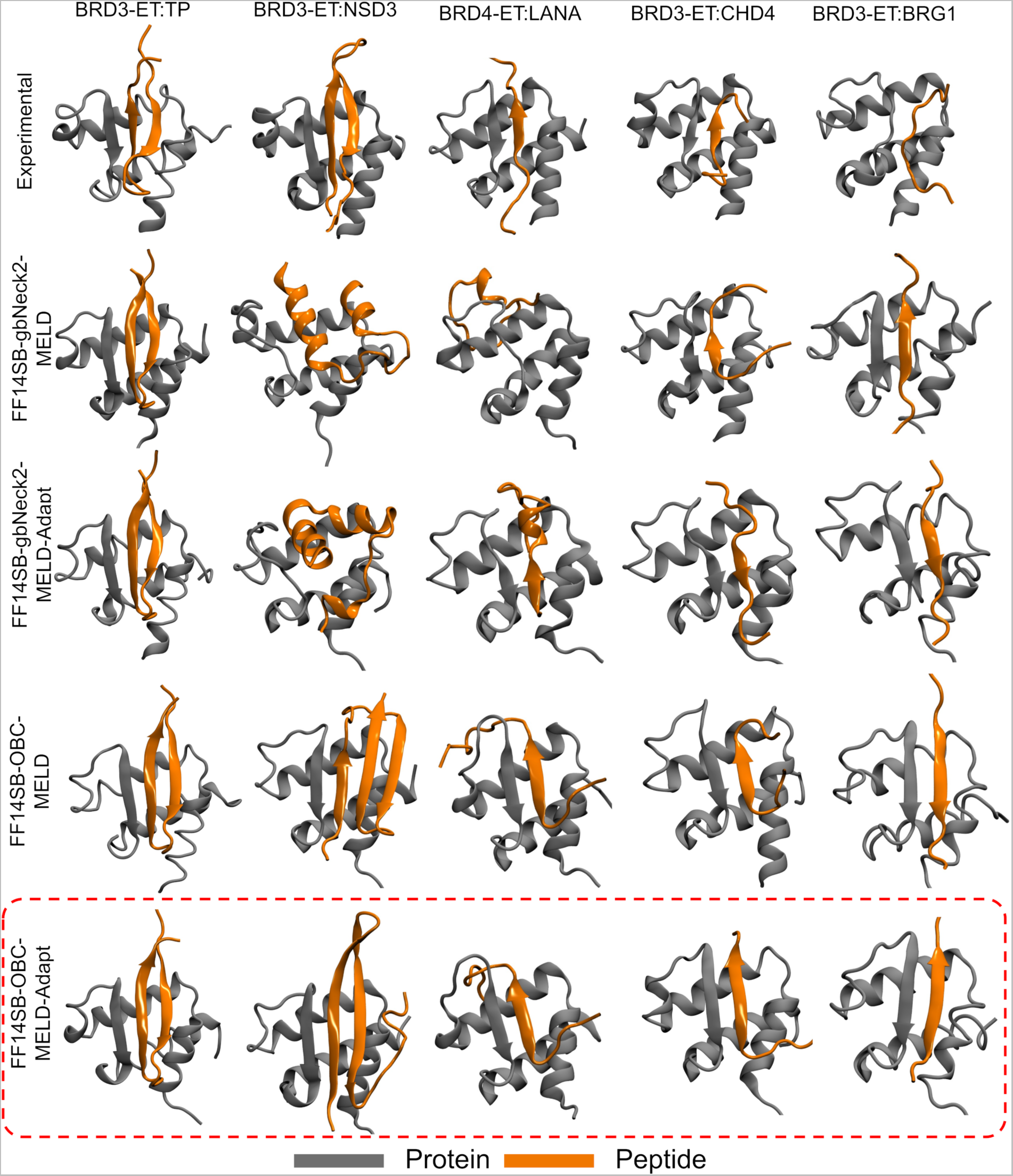
MELD-Adapt representative structures (centroid from the highest population cluster) for protein-peptide complexes simulated with different force fields. For each of the five simulated systems, the representative structure in each of the four simulation conditions are shown. The experimental structures are given as reference in the top row. The protein force field was fixed at the ff14SBside, with implicit solvent changing between OBC and GBneck2, and sampling using MELD or MELD-Adapt (λ = 0.25kcal/mol). The red dotted box shows the most succesful protocol.

Acknowledging that this could be a force field deficiency, rather than an issue with the MELD-Adapt procedure, we ran simulations with different force fields (see methods) to better balance preferences between the helical and extended conformations. ^34,35^ Indeed, in top predictions using the traditional MELD approach with the OBC implicit solvent and ff14SBside with CMAP^35^ correction significantly improved, capturing three of the five complexes as the top cluster (missing the orientation of BRG1 and the NSD3 peptide conformation). The results improved with MELD-Adapt simulations, with all five predictions corresponding to the native structure of the complex. Simulations with an earlier force field version (ff12SB) failed to reproduce most structures of the complex (even TP). Figures S17 and S18 help rationalize the balance between the force field and parameter sampling. For instance, all protocols and force fields were able to sample native-like structures of the complex (see RMSD values in Figure S18). They were just not preferred as high-population clusters in several cases. Figure S18 shows that the force field has a larger influence on the performance than whether MELD or MELD-Adapt was used. However, a head-to-head comparison between MELD and MELD-Adapt within each forcefield favors the later strategy. Overall pairs of systems, there was an insignificant improvement over the top five predictions and over the ensemble. However, the average MELD-Adapt improvement for the top cluster predictions was about 1-2 Å, showing that the approach is able to improve the ranking amongst force field selected top clusters, increasing its population.

### On-the-fly learning dataset accuracy improves modeling predictions

Reducing human intervention and decisions in computational modeling leads to greater reproducibility. In the context of integrative approaches, the user has many choices to make: from the number of relevant states, to how to model the data, to how to deal with uncertainties in the data. The MELD approach starts with the idea that if we have a belief in the data that is equal to or lower than the real accuracy of the data, it can identify the best interpretation of the data as well as the structures compatible with the data. However, estimating this value is not trivial, and will change according to experiments. The ability to sample native-like structures is still conditioned by challenges like backtracking,^36^ force field accuracy, and sampling efficiency – and it is hard to disambiguate the different contributions. Here, we have shown that on-the-fly identification of data accuracy can lead to more efficient simulations, increase sampling efficiency when our initial belief is too low, and correct phase space exploration when our initial belief drives to incorrect structures. However, the method is sensitive to the internal structure of the data (local or distributed) and the balance with the force field. Thus, high reward values can lead to too much data being satisfied and overcoming force field preferences. We did not find a protocol where fixing the reward value can be uniformly transferable across datasets, but lower reward values (0.25kcal/mol) seemed to work best overall in both cases – with low guiding power for the protein folding dataset.

## Conclusion

This study presents an enhanced MELD approach that incorporates on-the-fly learning to dynamically calibrate the use of experimental data, improving the accuracy of biomolecular structure predictions. Our results underline the importance of carefully balancing force field influences and data restraints to avoid convergence to incorrect regions of phase space. While the method shows sensitivity to the internal structure of the data and the balance with the force field, we have demonstrated that lower reward values tend to offer a safer compromise between data guidance and depending more on force field accuracy. The ability of the MELD-Adapt technique to recover from incorrect initial beliefs showcases its potential as a powerful tool in integrative structural biology, especially when addressing the inherent uncertainties present in experimental datasets. Ultimately, our approach reduces human intervention, increasing the reproducibility of computational modeling and providing a robust framework for predicting the structure of proteins and their complexes with peptides.

## Acknowledgement

AP thanks NSF CAREER CHE-2235785 for funding. JLM thanks XYZ for funding.

## Supporting Information Available

Supplementary tables and figures are provided.

