## Supplementary material for "MELD-Adapt: On-the-Fly Belief Updating in Integrative Molecular Dynamics": SI Tables and Figures

**Table S1.** Benchmark system used for protein folding using MELD-Adapt.

| PDB ID | Protein name |
|---|---|
| 3GB1 | Protein G |
| 1BDD | B domain of Staphylococcus Aureus Protein A |
| 1DV0 | C-terminal UBA domain of HHR23A |
| 1FEX | MYB-domain of human RAP1 |
| 1FME | Structure of FSD-EY |
| 1PRB | Albumin-binding domain |
| 2A3D | De novo designed three helix bundle protein |
| 2P6J | Designed engrailed homeodomain variant UVF |
| 2F4K | Villin subdomain HP-35 |
| 1UBQ | Ubiquitin |
| 2MQ8 (T0769) | De novo designed protein LFR1 1 with ferredoxin fold |
| 2N2U (T0773) | De novo designed ferredoxin fold protein SFR3 |

**Table S2.** First passage Time (ns) at Lowest Temperature Replica

| PDB ID | MELD | MELD-Adapt | | | | | |
|---|---|---|---|---|---|---|---|
| | | **0.25** | **0.5** | **1.0** | **2.0** | **4.0** | **8.0** |
| 3GB1 | 9.65 | 47.00 | 17.80 | 25.40 | 33.55 | 77.55 | none |
| 1BDD | 5.05 | 4.90 | 5.05 | 5.40 | 21.95 | 5.30 | 7.65 |
| 1DV0 | 4.85 | 4.35 | 3.30 | 5.20 | 6.30 | 10.30 | 4.65 |
| 1FEX | 10.70 | 12.05 | 5.85 | 30.70 | 24.50 | none | none |
| 1FME | 7.70 | 13.95 | 13.95 | 8.95 | 11.95 | 10.85 | 8.45 |
| 1PRB | 15.2 | 11.7 | 12.10 | 13.95 | 33.55 | none | none |
| 2A3D | 5.00 | 9.05 | 6.00 | 9.20 | 18.50 | none | none |
| 2P6J | 6.40 | 5.80 | 6.80 | 5.20 | 3.55 | 3.40 | 5.30 |
| 2F4K | 5.65 | 22.75 | 9.15 | 14.90 | 59.45 | 21.45 | none |
| 1UBQ | 72.15 | 391.25 | 232.65 | 210.05 | 20.00 | 84.30 | none |

**Table S3**. RMSD/ Å of best structure, Top1 cluster and Top 5 cluster between MELD and MELD-Adapt. The value in parenthesis refers to the cluster number.

| PDB ID | | MELD | | | MELD-Adapt (reward = 1.0 kcal/mol) | | |
|---|---|---|---|---|---|---|---|
| | | **best** | **top1** | **top5** | **best** | **top1** | **top5** |
| 3GB1 | underestimate | 1.11 | 1.22 | 1.22 (c1) | 1.11 | 1.33 | 1.33 (c1) |
| | correct estimate | 1.13 | 1.30 | 1.30 (c1) | 1.11 | 1.28 | 1.28 (C1) |
| | overestimate | 5.80 | 9.89 | 8.31 (c2) | 1.11 | 1.28 | 1.28 (c1) |
| T0769 | underestimate | 1.70 | 2.09 | 2.09 (c1) | 1.62 | 2.44 | 2.44 (c1) |
| | correct estimate | 1.64 | 2.74 | 2.46 (c5) | 1.88 | 3.16 | 2.73 (c4) |
| | overestimate | 1.60 | 2.16 | 2.16 (c1) | 2.14 | 3.25 | 2.52 (c2) |
| T0773 | underestimate | 1.30 | 4.88 | 4.88(c1) | 1.31 | 1.79 | 1.79(c1) |
| | correct estimate | 1.34 | 1.81 | 1.81 (c1) | 1.42 | 2.06 | 2.06 (c1) |
| | overestimate | 1.34 | 2.00 | 2.00 (c1) | 1.39 | 1.93 | 1.93 (c1) |

**Table S4**. Percentage Population of Top1 cluster between MELD and MELD-Adapt.

| PDB ID | | MELD | MELD-Adapt (reward = 1.0 kcal/mol) |
|---|---|---|---|
| 3GB1 | underestimate | 64.4 | 97.8 |
| | correct estimate | 77.6 | 100 |
| | overestimate | 18.4 | 98.1 |
| T0769 | underestimate | 13.9 | 100 |
| | correct estimate | 94.8 | 99.9 |
| | overestimate | 99.8 | 99.9 |
| T0773 | underestimate | 14.1 | 69.2 |
| | correct estimate | 98.2 | 72.1 |
| | overestimate | 97.6 | 44.7 |

**Table S5**. Comparison of First passage Time (ns) at Lowest Temperature Replica between MELD and MELD-Adapt.

| PDB ID | | MELD | MELD-Adapt (reward = 1.0 kcal/mol) |
|---|---|---|---|
| 3GB1 | underestimate | 122.35 | 10.30 |
| | correct estimate | 9.65 | 25.40 |
| | overestimate | none | 87.55 |
| T0769 | underestimate | none | 23.65 |
| | correct estimate | 21.45 | 4.00 |
| | overestimate | 10.05 | 4.55 |
| T0773 | underestimate | 28.50 | 19.30 |
| | correct estimate | 22.55 | 33.45 |
| | overestimate | 13.30 | 110.65 |

**Table S6**. Benchmark system used for protein-peptide binding using MELD-Adapt.

| PDB ID | System name (protein:peptide) | Length (protein:peptide) | Binding affinity |
|---|---|---|---|
| 7JQ8 | BRD3-ET:TP | 68:23 | 90 nM |
| 6BGH | BRD3-ET:BRG1 | 68:12 | 7 μM |
| 6BGG | BRD3-ET:CHD4 | 68:12 | 95 μM |
| 7JYN | BRD3-ET:NSD3 | 68:34 | 250 μM |
| 2ND0 | BRD4-ET:LANA | 68:19 | 635 μM |

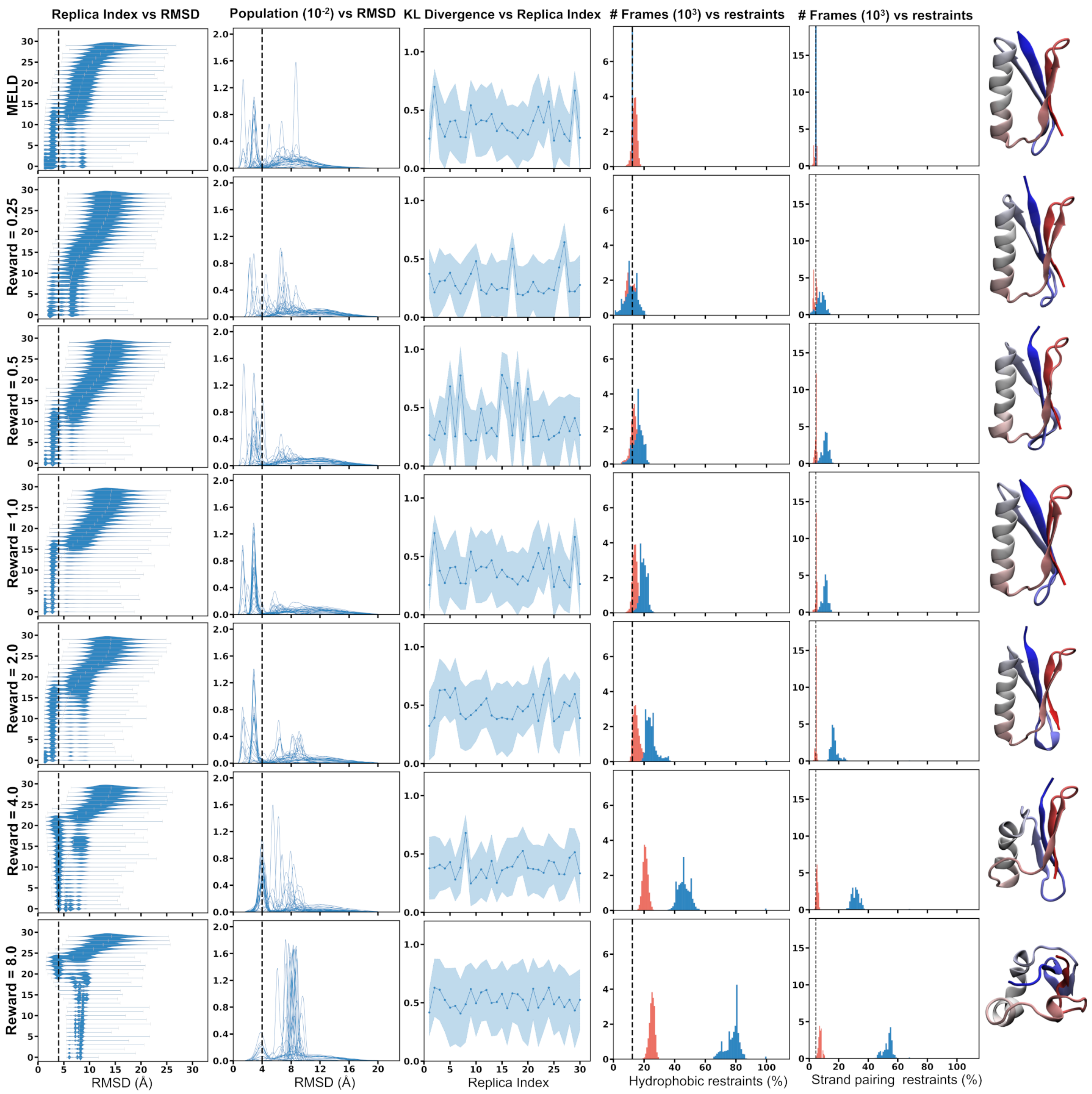


**Figure S1. Comparative analysis of protocols with varied reward values for 3GB1.** Each panel illustrates the outcomes of protocols with fixed parameters and reward values of 0.25, 0.5, 1.0, 2.0, 4.0, and 8.0. The header in each column provides axis titles for clarity. The panels from left to right include: the funneling protocol (described in the Main text), RMSD distributions across 30 walkers relative to the native structure, average pair wise KL divergence over the 30 walkers, the percentage of active restraints during simulation (blue) and the percentage of satisfied hydrophobic restraints (red), the percentage of active restraints during simulation (blue) and the percentage of satisfied strand pairing restraints (red). Dotted vertical lines in the first and second columns represent an RMSD cutoff of 4Å, indicating a significant deviation from the native structure. Additionally, dotted vertical lines in the fourth and fifth columns denote the number of restraints satisfied in the native structure. The sixth column showcases the Top cluster representative structure.

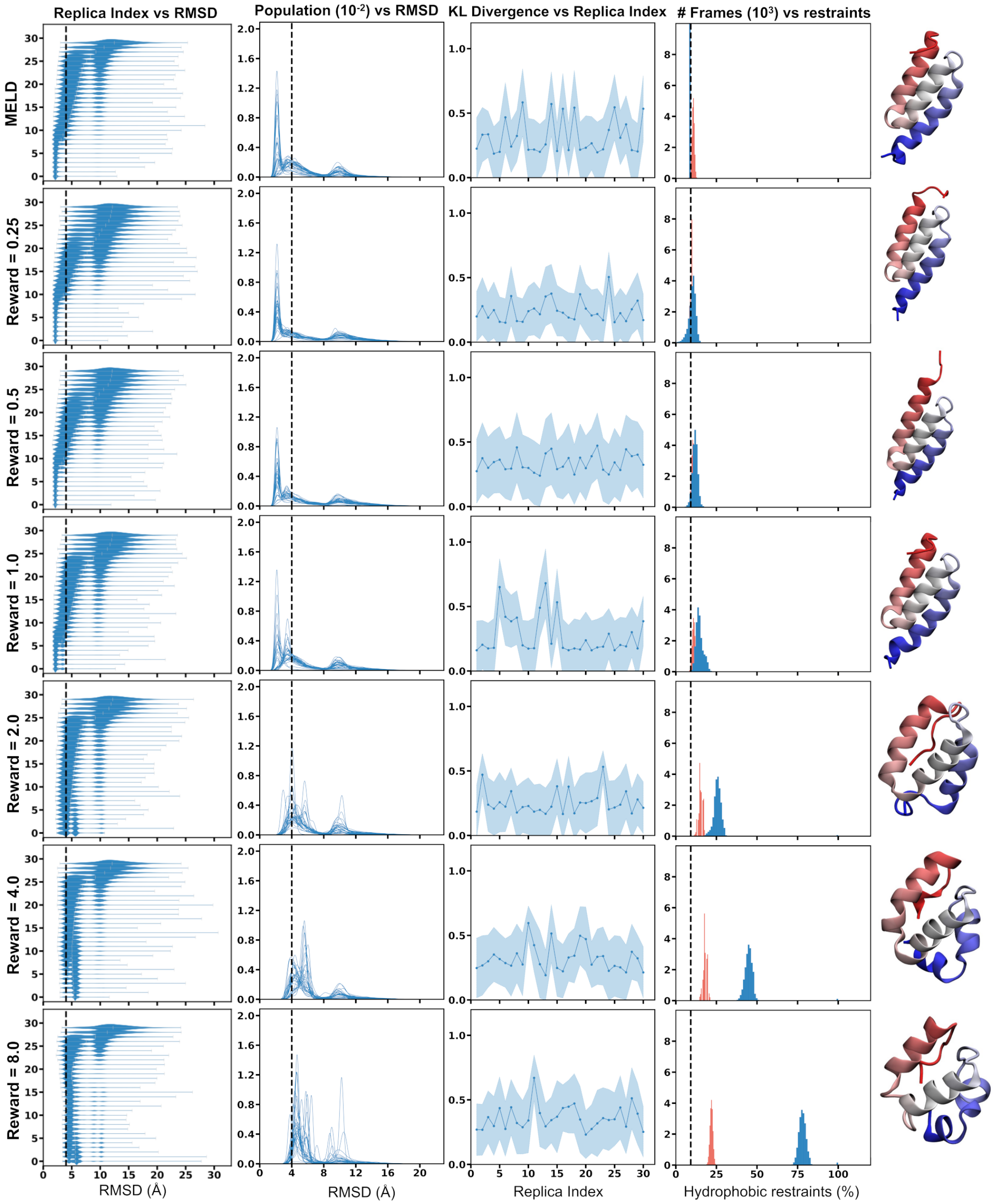


**Figure S2. Comparative analysis of protocols with varied reward values for 1BDD.** Each panel illustrates the outcomes of protocols with fixed parameters and reward values of 0.25, 0.5, 1.0, 2.0, 4.0, and 8.0. The header in each column provides axis titles for clarity. The panels from left to right include: the funneling protocol (described in the Main text), RMSD distributions across 30 walkers relative to the native structure, average pair wise KL divergence over the 30 walkers, the percentage of active restraints during simulation (in blue) and the percentage of satisfied hydrophobic restraints (in red). Dotted vertical lines in the first and second columns represent an RMSD cutoff of 4Å, indicating a significant deviation from the native structure. Additionally, dotted vertical lines in the fourth column denotes the number of restraints satisfied in the native structure. The fifth column showcases the Top cluster representative structure.

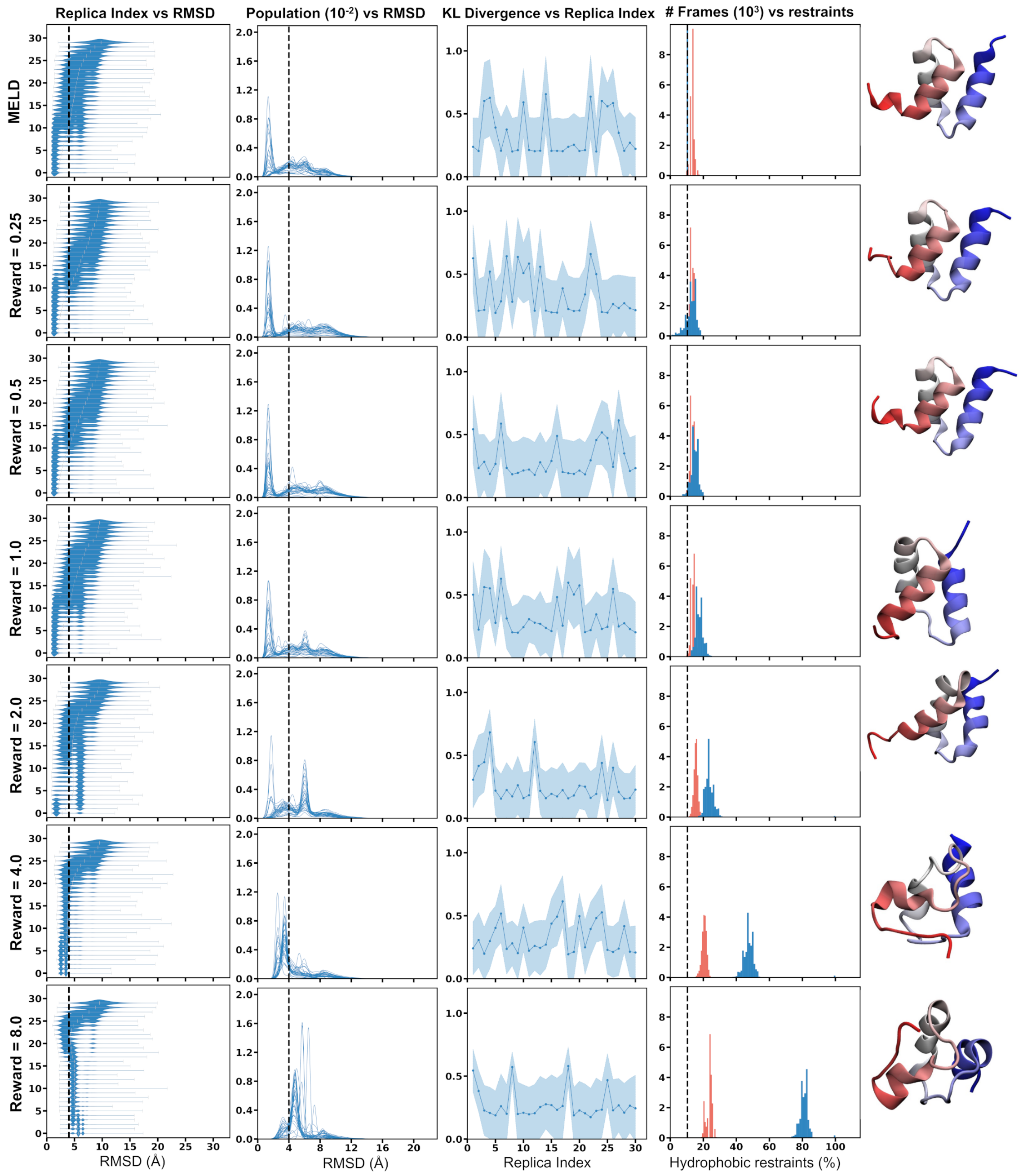


**Figure S3. Comparative analysis of protocols with varied reward values for 1DV0.** Each panel illustrates the outcomes of protocols with fixed parameters and reward values of 0.25, 0.5, 1.0, 2.0, 4.0, and 8.0. The header in each column provides axis titles for clarity. The panels from left to right include: the funneling protocol (described in the Main text), RMSD distributions across 30 walkers relative to the native structure, average pair wise KL divergence over the 30 walkers, the percentage of active restraints during simulation (in blue) and the percentage of satisfied hydrophobic restraints (in red). Dotted vertical lines in the first and second columns represent an RMSD cutoff of 4Å, indicating a significant deviation from the native structure. Additionally, dotted vertical lines in the fourth column denotes the number of restraints satisfied in the native structure. The fifth column showcases the Top cluster representative structure.

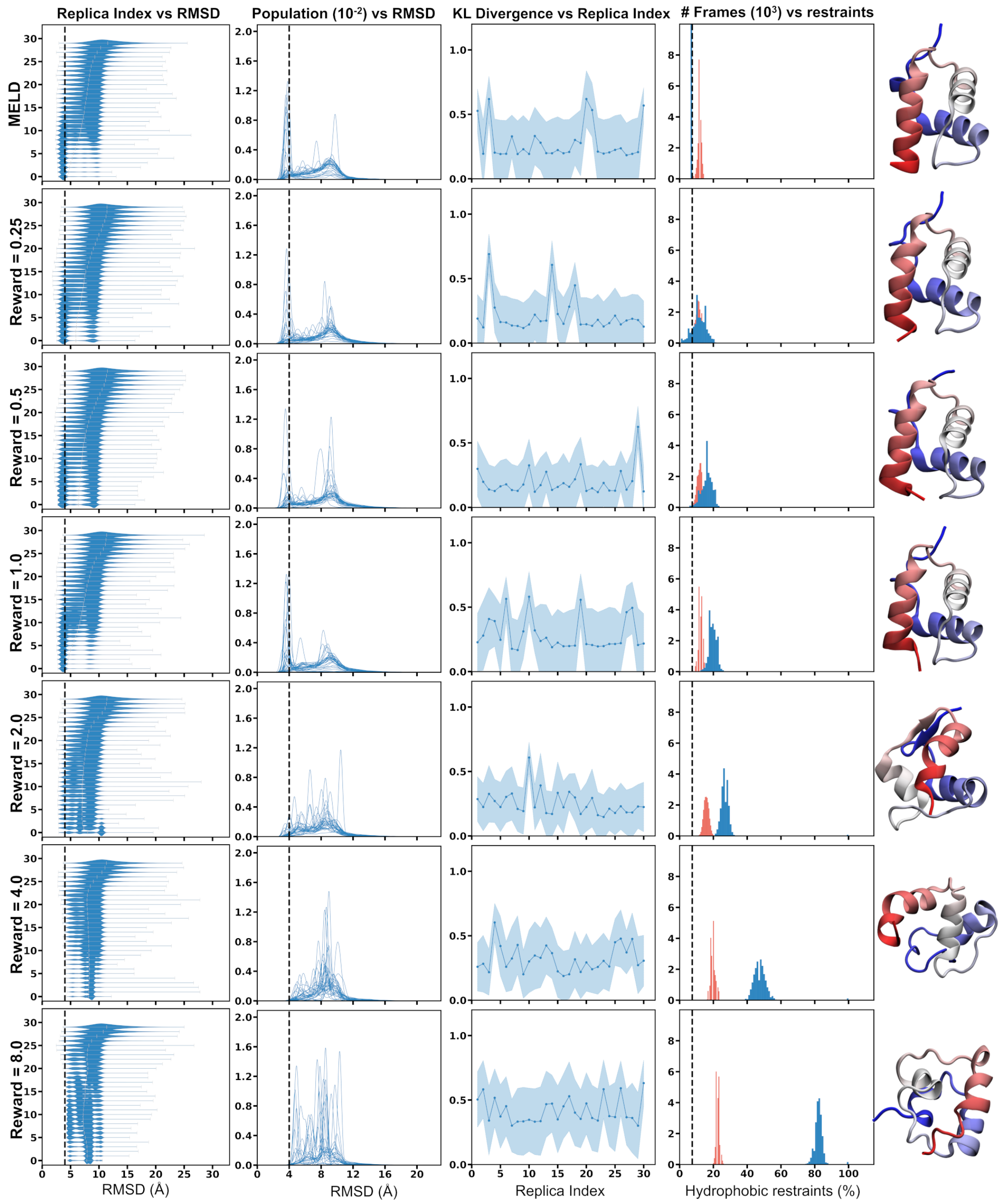


**Figure S4. Comparative analysis of protocols with varied reward values for 1FEX.** Each panel illustrates the outcomes of protocols with fixed parameters and reward values of 0.25, 0.5, 1.0, 2.0, 4.0, and 8.0. The header in each column provides axis titles for clarity. The panels from left to right include: the funneling protocol (described in the Main text), RMSD distributions across 30 walkers relative to the native structure, average pair wise KL divergence over the 30 walkers, the percentage of active restraints during simulation (in blue) and the percentage of satisfied hydrophobic restraints (in red). Dotted vertical lines in the first and second columns represent an RMSD cutoff of 4Å, indicating a significant deviation from the native structure. Additionally, dotted vertical lines in the fourth column denotes the number of restraints satisfied in the native structure. The fifth column showcases the Top cluster representative structure.

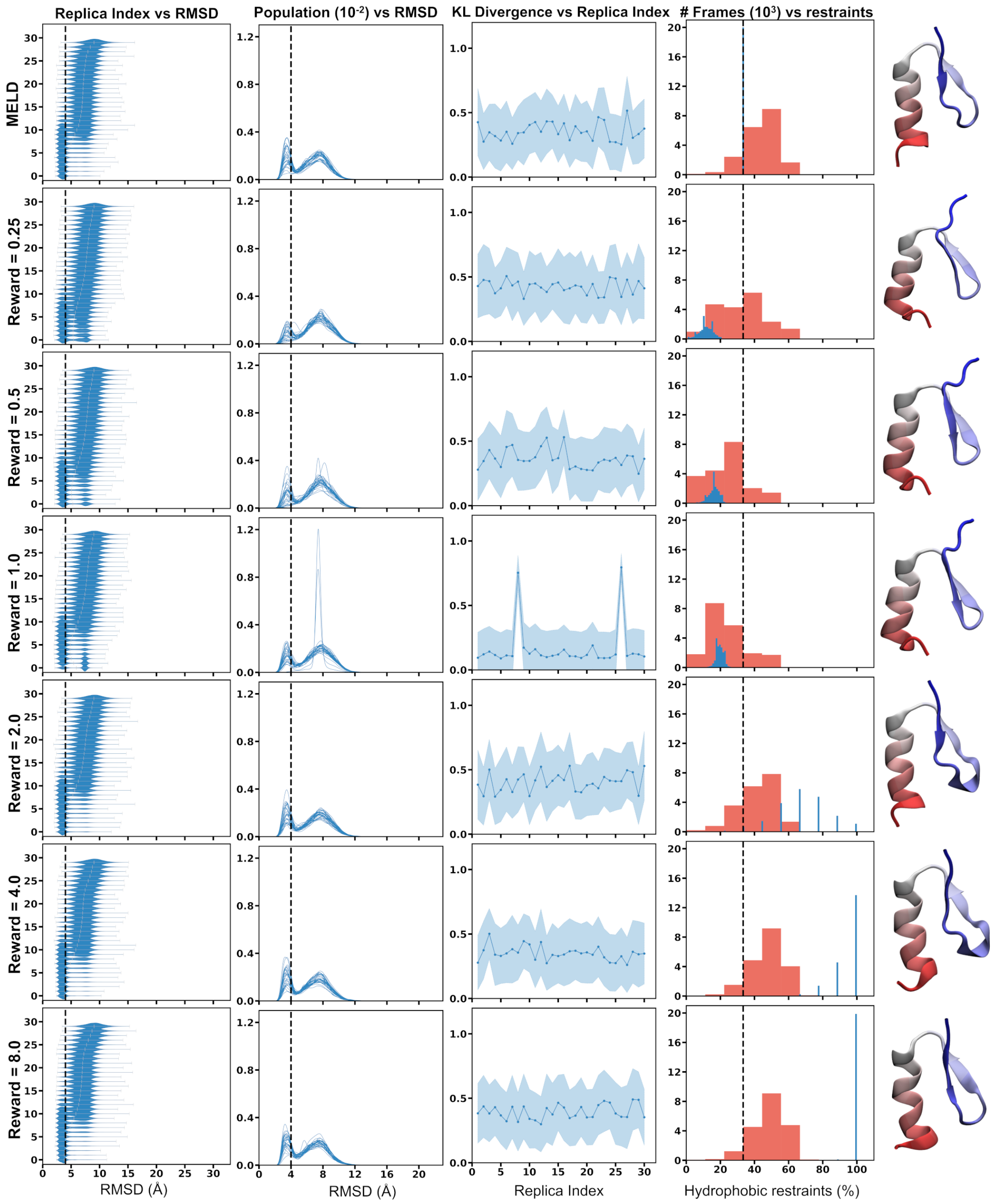


**Figure S5. Comparative analysis of protocols with varied reward values for 1FME.** Each panel illustrates the outcomes of protocols with fixed parameters and reward values of 0.25, 0.5, 1.0, 2.0, 4.0, and 8.0. The header in each column provides axis titles for clarity. The panels from left to right include: the funneling protocol (described in the Main text), RMSD distributions across 30 walkers relative to the native structure, average pair wise KL divergence over the 30 walkers, the percentage of active restraints during simulation (in blue) and the percentage of satisfied hydrophobic restraints (in red). Dotted vertical lines in the first and second columns represent an RMSD cutoff of 4Å, indicating a significant deviation from the native structure. Additionally, dotted vertical lines in the fourth column denotes the number of restraints satisfied in the native structure. The fifth column showcases the Top cluster representative structure.

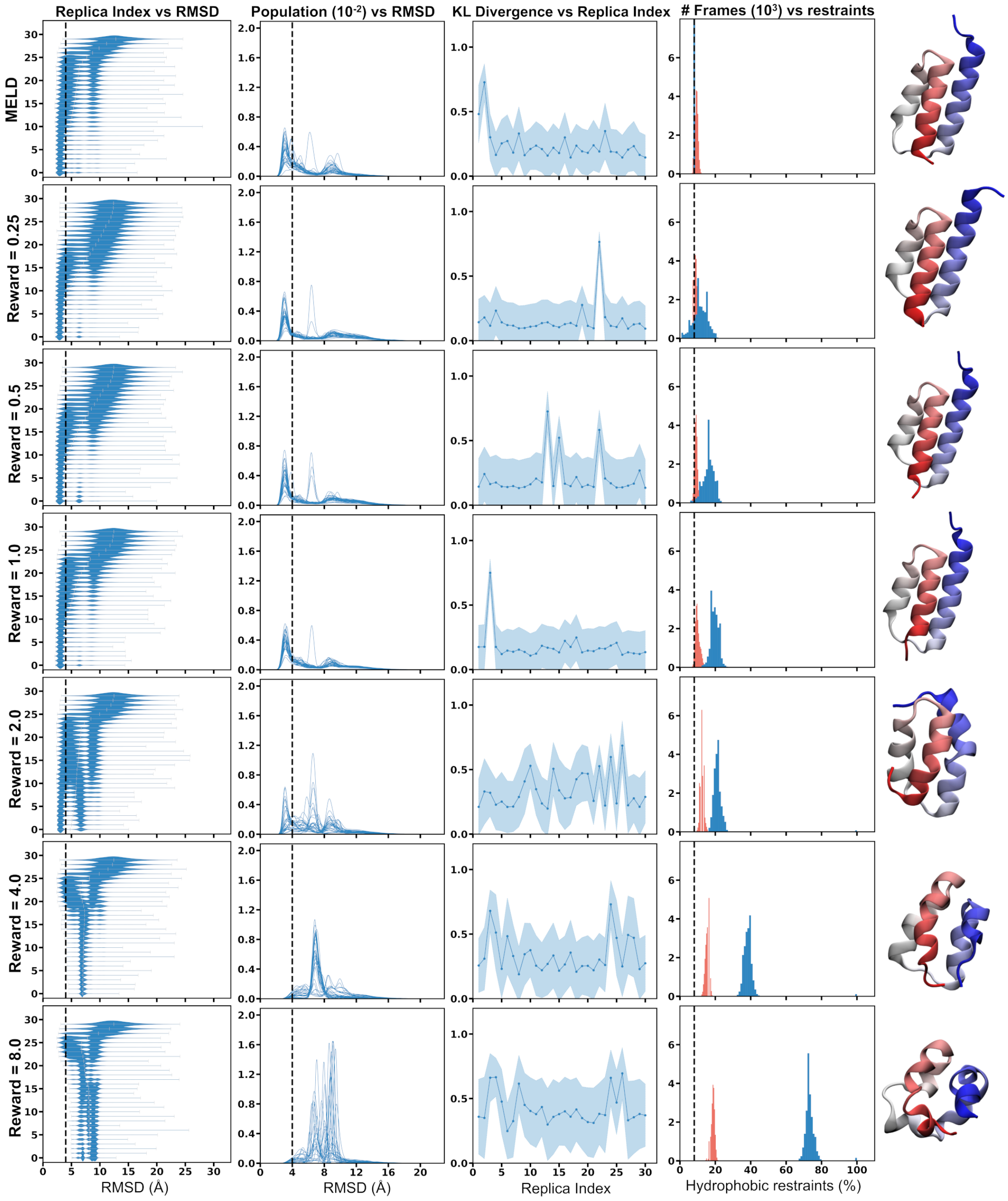


**Figure S6. Comparative analysis of protocols with varied reward values for 1PRB.** Each panel illustrates the outcomes of protocols with fixed parameters and reward values of 0.25, 0.5, 1.0, 2.0, 4.0, and 8.0. The header in each column provides axis titles for clarity. The panels from left to right include: the funneling protocol (described in the Main text), RMSD distributions across 30 walkers relative to the native structure, average pair wise KL divergence over the 30 walkers, the percentage of active restraints during simulation (in blue) and the percentage of satisfied hydrophobic restraints (in red). Dotted vertical lines in the first and second columns represent an RMSD cutoff of 4Å, indicating a significant deviation from the native structure. Additionally, dotted vertical lines in the fourth column denotes the number of restraints satisfied in the native structure. The fifth column showcases the Top cluster representative structure.

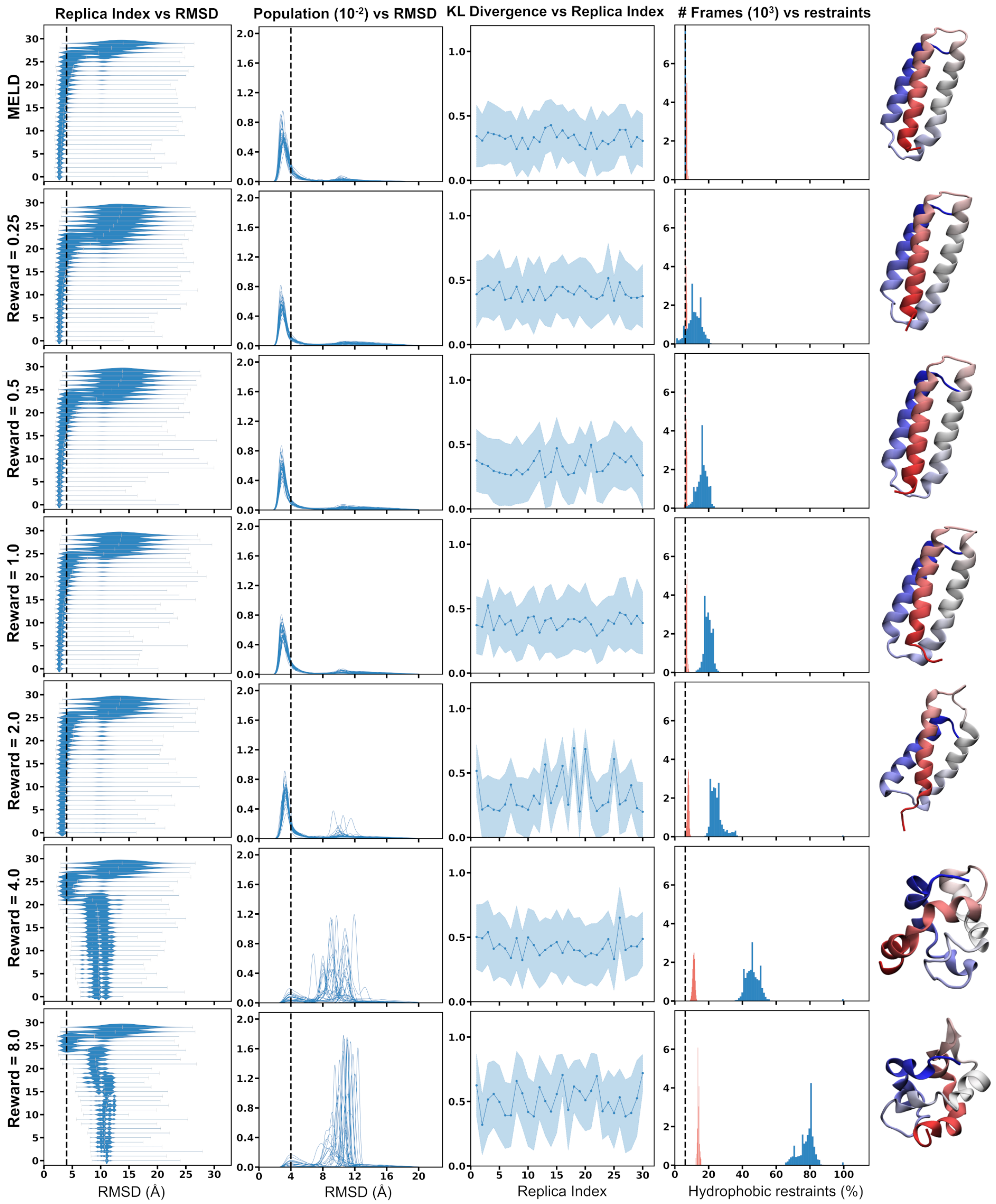


**Figure S7. Comparative analysis of protocols with varied reward values for 2A3D.** Each panel illustrates the outcomes of protocols with fixed parameters and reward values of 0.25, 0.5, 1.0, 2.0, 4.0, and 8.0. The header in each column provides axis titles for clarity. The panels from left to right include: the funneling protocol (described in the Main text), RMSD distributions across 30 walkers relative to the native structure, average pair wise KL divergence over the 30 walkers, the percentage of active restraints during simulation (in blue) and the percentage of satisfied hydrophobic restraints (in red). Dotted vertical lines in the first and second columns represent an RMSD cutoff of 4Å, indicating a significant deviation from the native structure. Additionally, dotted vertical lines in the fourth column denotes the number of restraints satisfied in the native structure. The fifth column showcases the Top cluster representative structure.

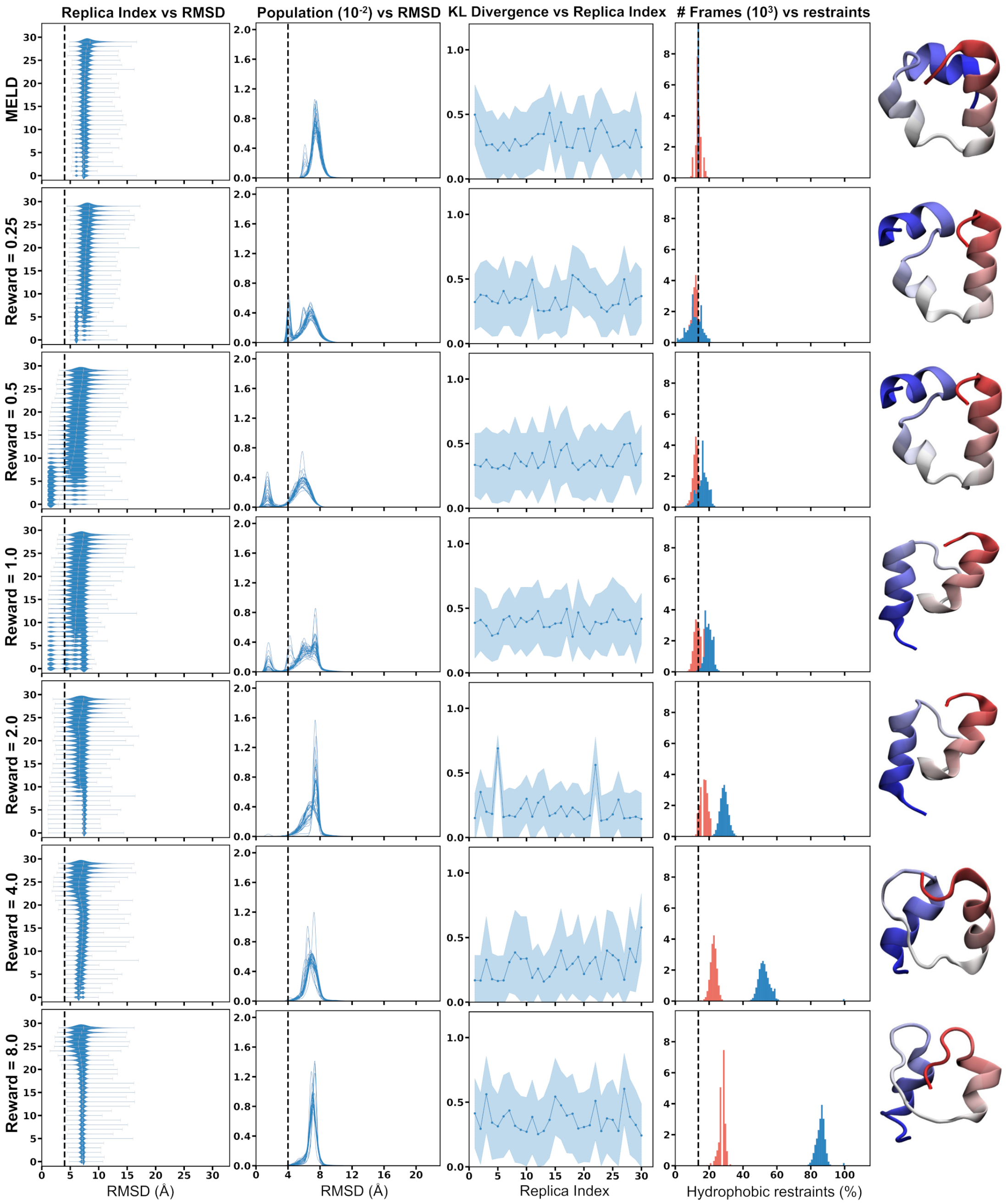


**Figure S8. Comparative analysis of protocols with varied reward values for 2F4K.** Each panel illustrates the outcomes of protocols with fixed parameters and reward values of 0.25, 0.5, 1.0, 2.0, 4.0, and 8.0. The header in each column provides axis titles for clarity. The panels from left to right include: the funneling protocol (described in the Main text), RMSD distributions across 30 walkers relative to the native structure, average pair wise KL divergence over the 30 walkers, the percentage of active restraints during simulation (in blue) and the percentage of satisfied hydrophobic restraints (in red). Dotted vertical lines in the first and second columns represent an RMSD cutoff of 4Å, indicating a significant deviation from the native structure. Additionally, dotted vertical lines in the fourth column denotes the number of restraints satisfied in the native structure. The fifth column showcases the Top cluster representative structure.

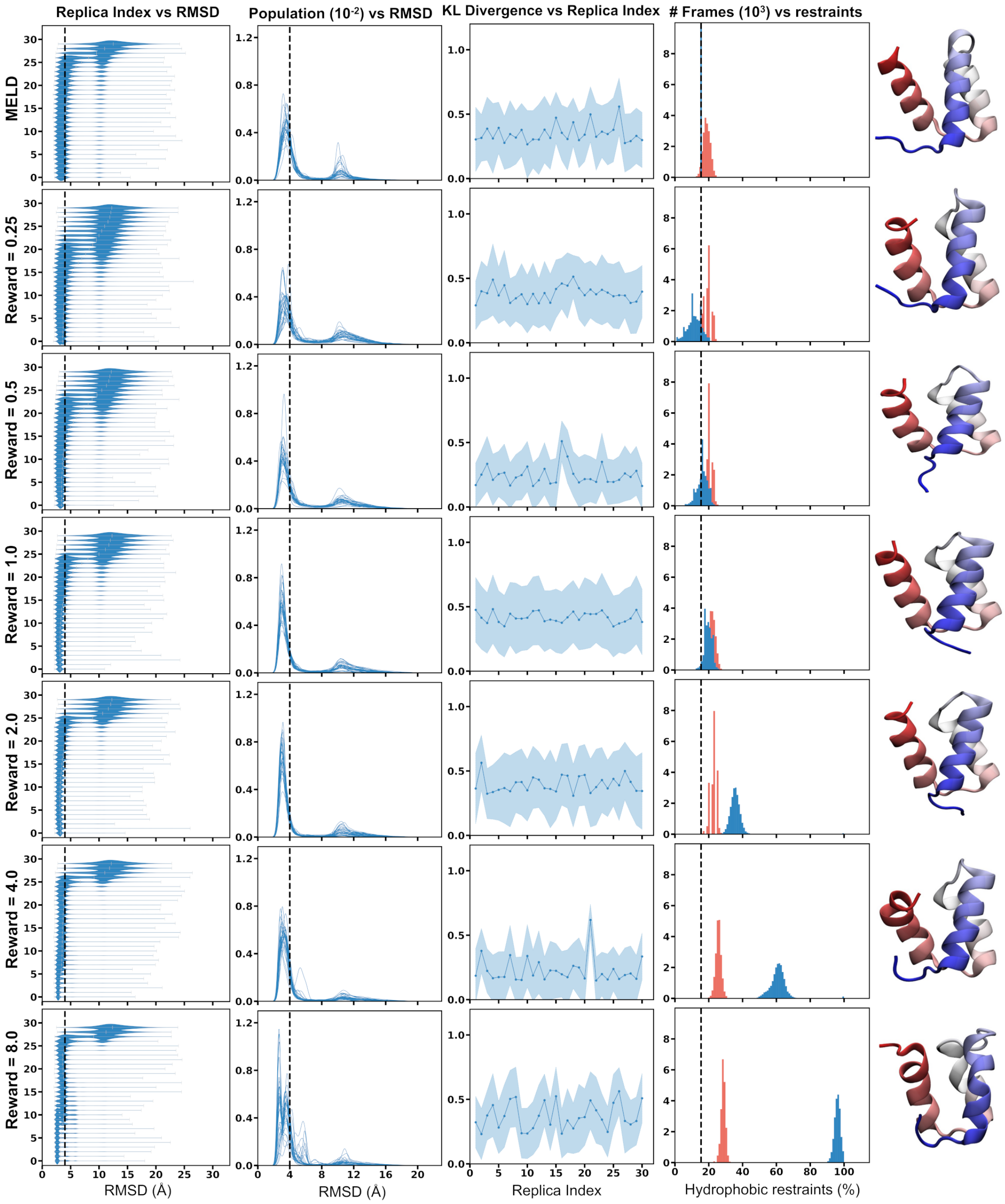


**Figure S9. Comparative analysis of protocols with varied reward values for 2P6J.** Each panel illustrates the outcomes of protocols with fixed parameters and reward values of 0.25, 0.5, 1.0, 2.0, 4.0, and 8.0. The header in each column provides axis titles for clarity. The panels from left to right include: the funneling protocol (described in the Main text), RMSD distributions across 30 walkers relative to the native structure, average pair wise KL divergence over the 30 walkers, the percentage of active restraints during simulation (in blue) and the percentage of satisfied hydrophobic restraints (in red). Dotted vertical lines in the first and second columns represent an RMSD cutoff of 4Å, indicating a significant deviation from the native structure. Additionally, dotted vertical lines in the fourth column denotes the number of restraints satisfied in the native structure. The fifth column showcases the Top cluster representative structure.

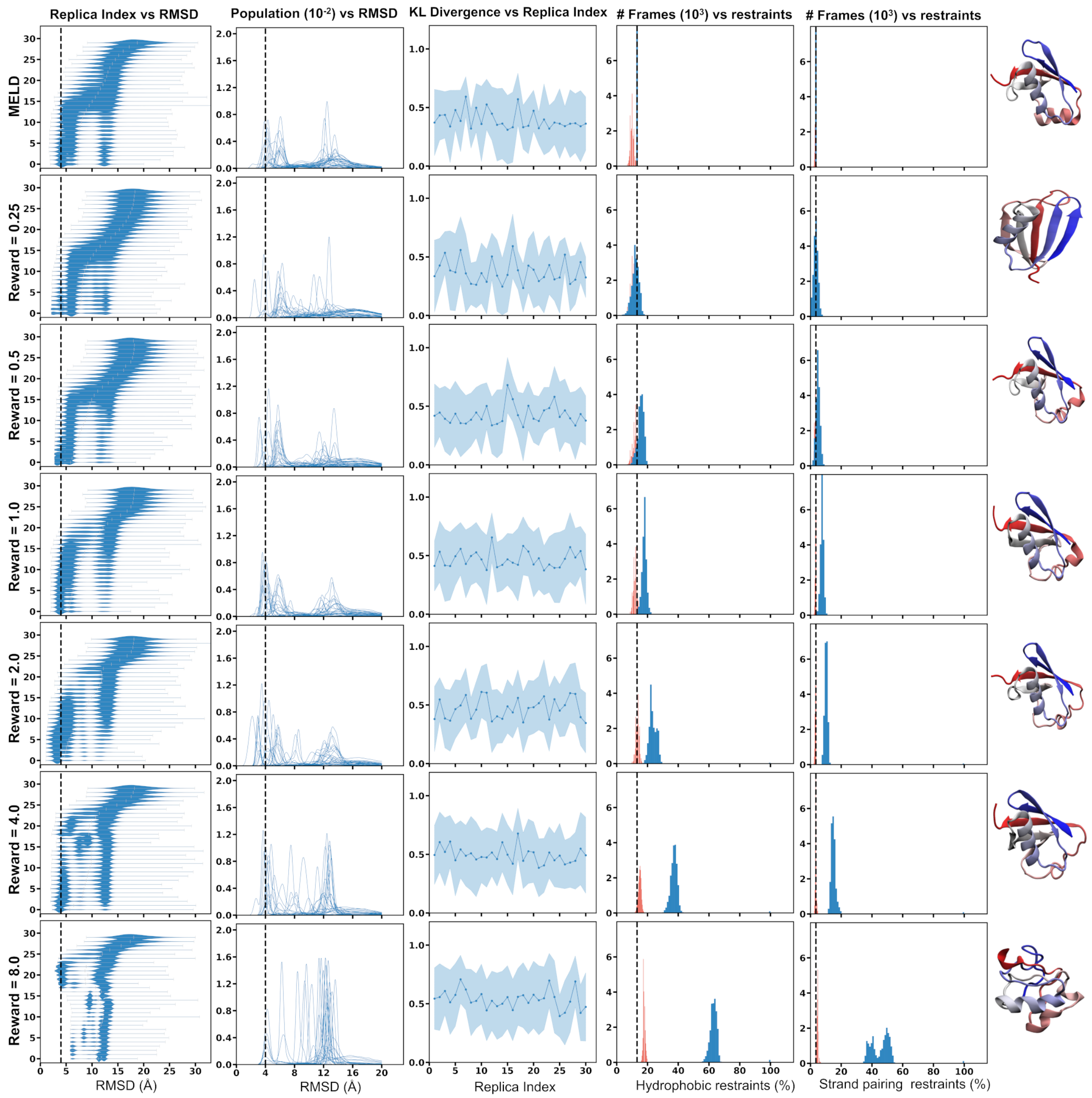


**Figure S10. Comparative analysis of protocols with varied reward values for 1UBQ.** Each panel illustrates the outcomes of protocols with fixed parameters and reward values of 0.25, 0.5, 1.0, 2.0, 4.0, and 8.0. The header in each column provides axis titles for clarity. The panels from left to right include: the funneling protocol (described in the Main text), RMSD distributions across 30 walkers relative to the native structure, average pair wise KL divergence over the 30 walkers, the percentage of active restraints during simulation (blue) and the percentage of satisfied hydrophobic restraints (red), the percentage of active restraints during simulation (blue) and the percentage of satisfied strand pairing restraints (red). Dotted vertical lines in the first and second columns represent an RMSD cutoff of 4Å, indicating a significant deviation from the native structure. Additionally, dotted vertical lines in the fourth and fifth columns denote the number of restraints satisfied in the native structure. The sixth column showcases the Top cluster representative structure.

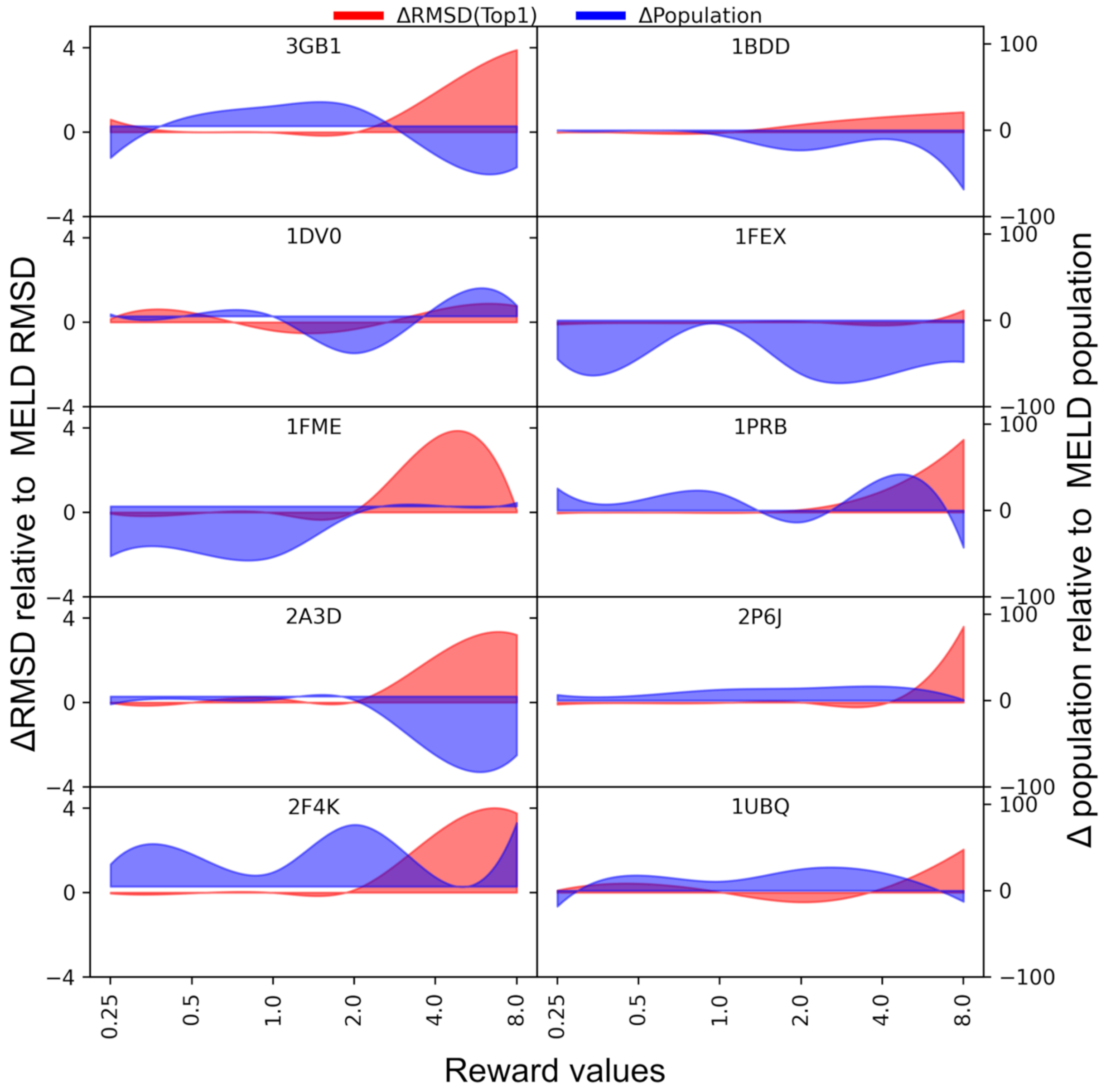


**Figure S11.** Differences in root mean square deviation (RMSD) of representative structures within the top cluster are compared across different reward values of MELD-Adapt relative to MELD (filled in red), alongside percentage differences in the population of the top cluster structure (secondary y-axis, filled in blue). Negative values indicate better reward values compared to MELD for RMSD difference (red), while higher positive values indicate better reward values compared to MELD for population difference (blue). These comparisons are illustrated for all 10 protein systems used as benchmarks for protein folding.

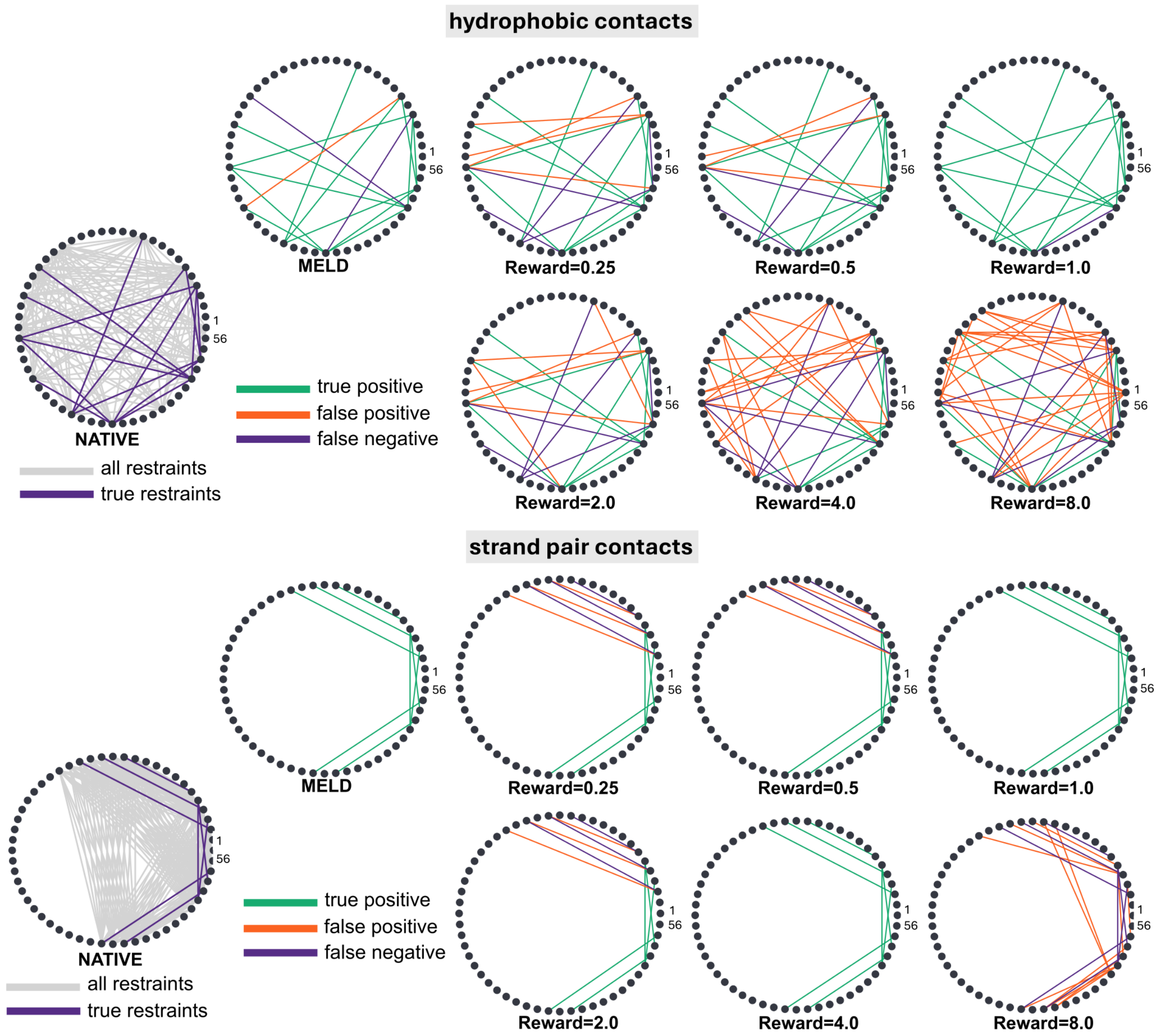


**Figure S12. Circular plots depicting the combinatorial possibilities of interactions between residues of protein 3GB1.** Each line connecting pairs of residues signifies a potential contact. Grey lines denote all possible contacts. Purple lines highlight true contacts in the native structure but absent in the predicted structure. Green lines indicate true positives (contacts present in both predicted and native structures), while red lines denote false positives (contacts present only in predicted structures). The top panel illustrates hydrophobic restraints, while the bottom panel illustrates strand pairing restraints.

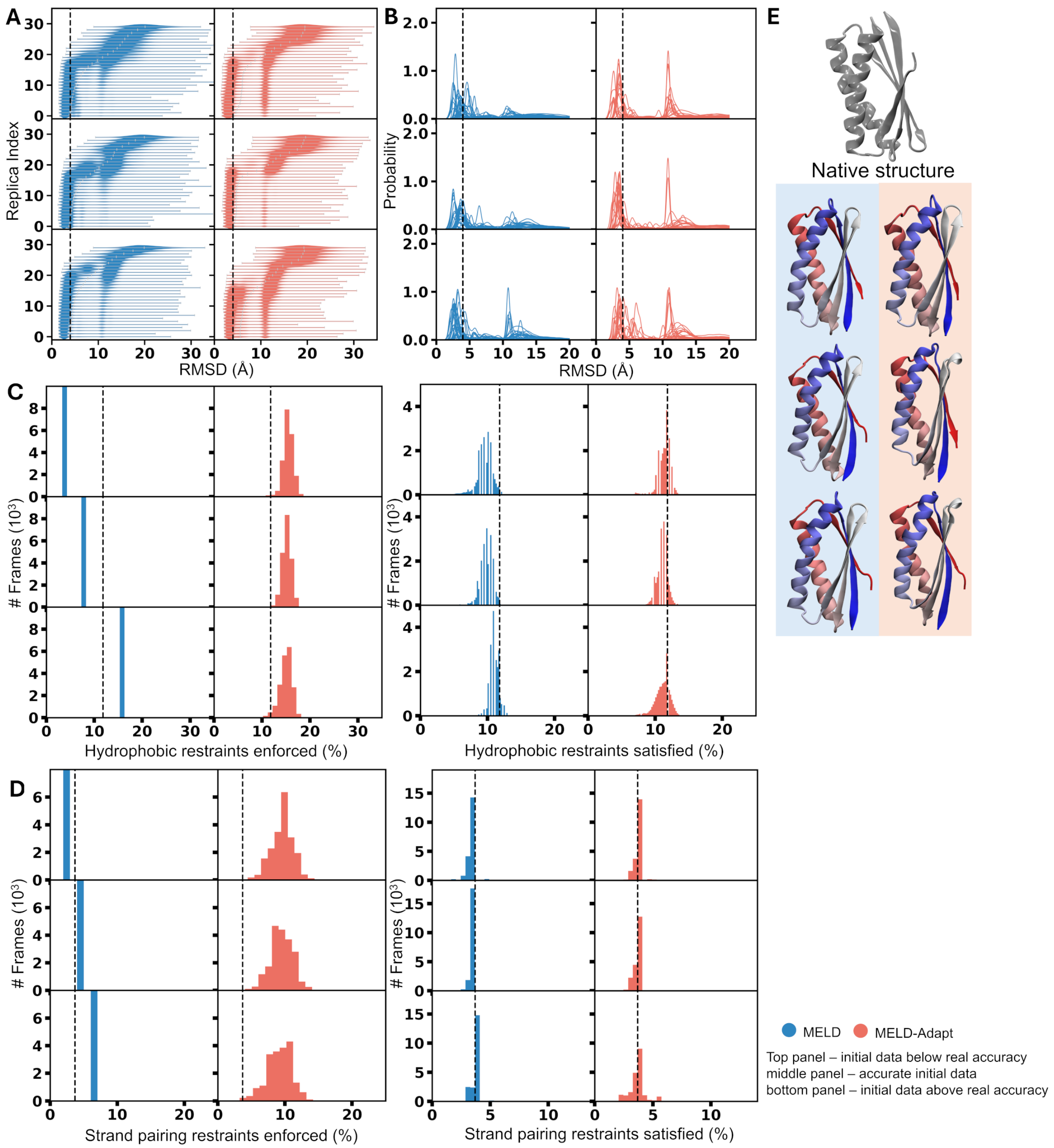


**Figure S13. Folding Belief Analysis for 2MQ8 (T0769).** Comparison of protein folding dynamics between MELD (blue) and MELD-Adapt (red, with a reward value of 1.0 kcal/mol) under three scenarios: i) initial data below real accuracy, ii) accurate initial data, and iii) initial data above real accuracy. The analysis includes A) Funneling protocol, B) RMSD distribution, C) percentage of enforced and satisfied hydrophobic restraints, D) percentage of enforced and satisfied strand pairing restraints, illustrating MELD-Adapt dynamically adjust the trust percentage and is independent from initial beliefs. E) Best predicted structures in each protocol compared to the native structure (silver).

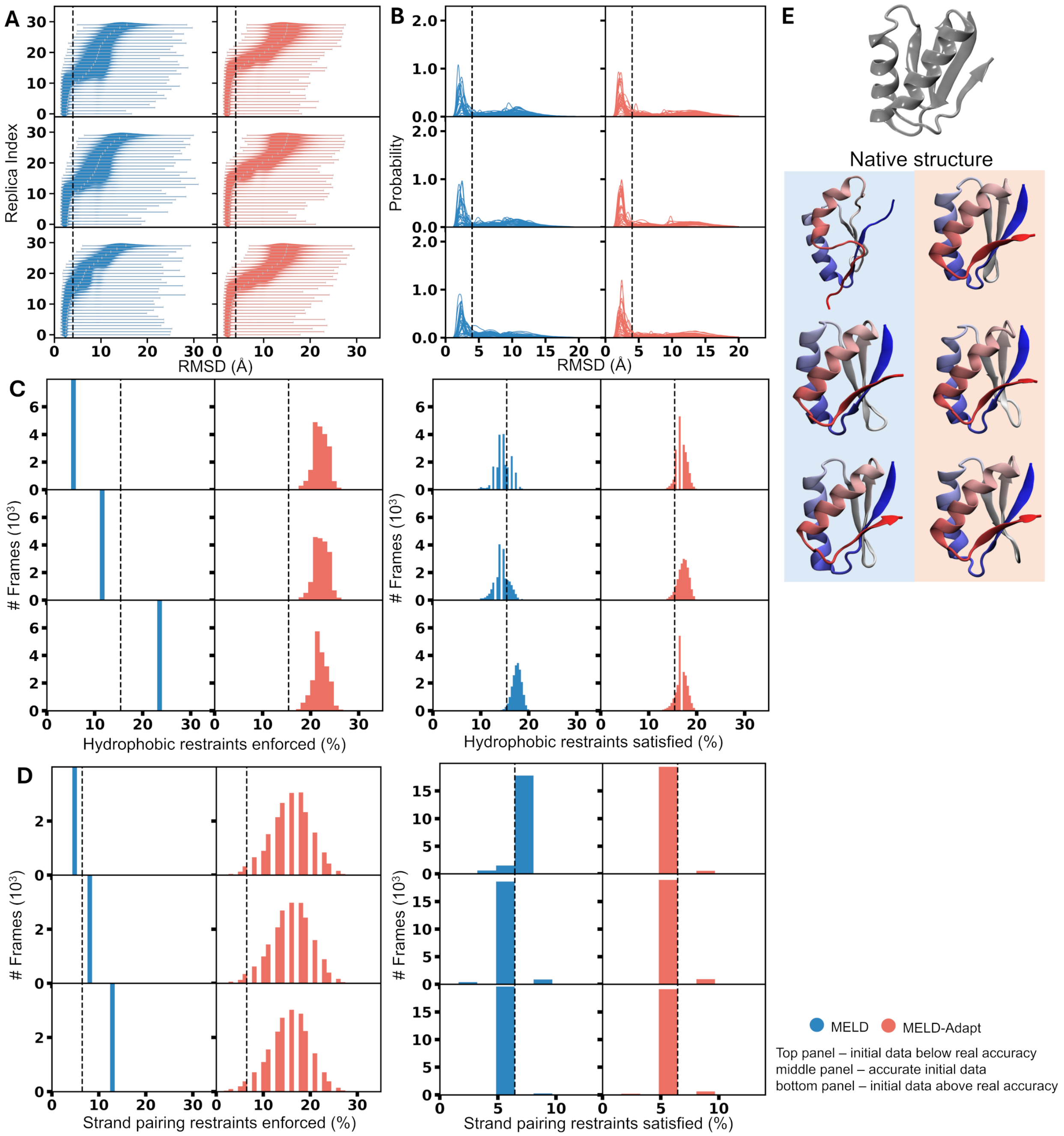


**Figure S14. Folding Belief Analysis for 2N2U (T0773).** Comparison of protein folding dynamics between MELD (blue) and MELD-Adapt (red, with a reward value of 1.0 kcal/mol) under three scenarios: i) initial data below real accuracy, ii) accurate initial data, and iii) initial data above real accuracy. The analysis includes A) Funneling protocol, B) RMSD distribution, C) percentage of enforced and satisfied hydrophobic restraints, D) percentage of enforced and satisfied strand pairing restraints, illustrating MELD-Adapt dynamically adjust the trust percentage and is independent from initial beliefs. E) Best predicted structures in each protocol compared to the native structure (silver).

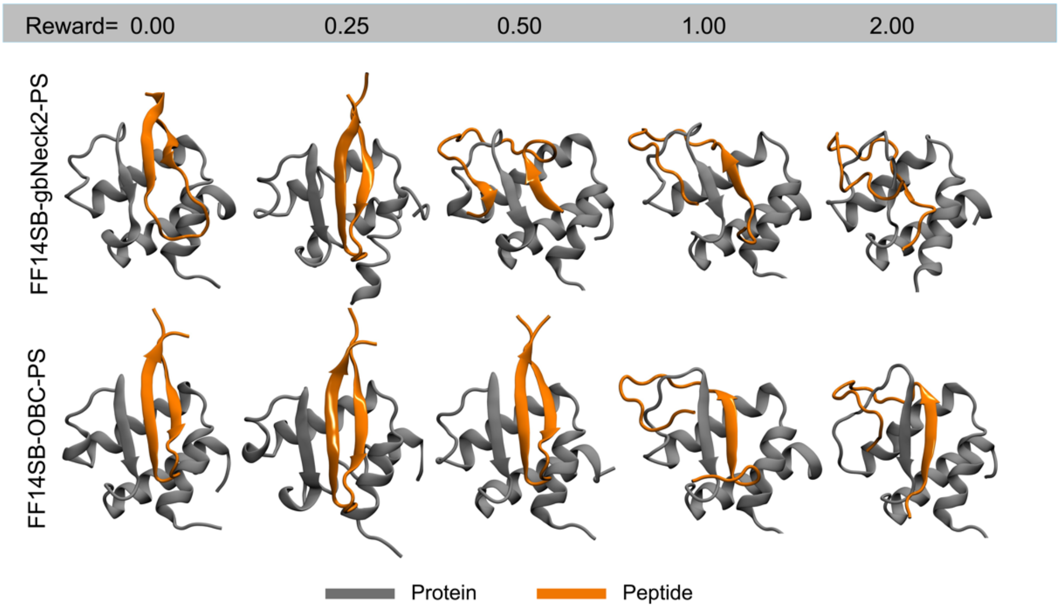


**Figure S15. Optimization of the reward values for protein-peptide binding.** Centroid of the highest populated cluster for BRD3-TP binding simulation at different reward values. The top row represents results from the simulations with FF14SBside force field with gbNeck2 implicit model, whereas the bottom row represents the results from simulations with OBC implicit model pair with the same force field.

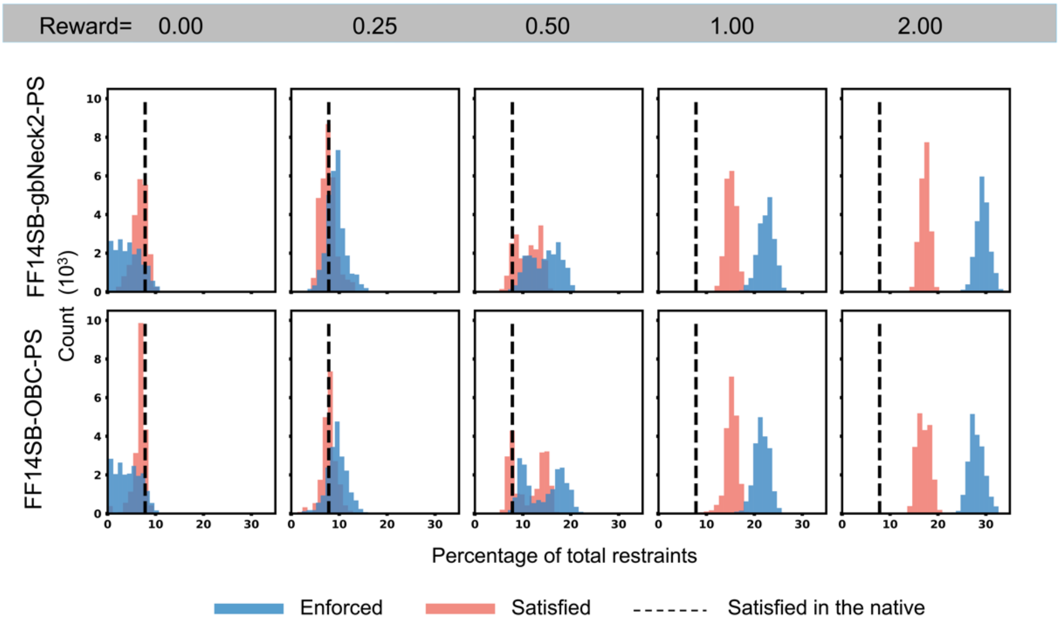


**Figure S16. Restraint distributions at different reward values for protein-peptide binding.** The distribution of enforced and satisfied contacts in the lowest temperature replica is plotted here along with the number of satisfied contacts in the experimental structure. The top row represents results from the simulations with FF14SBside force field with gbNeck2 implicit model, whereas the bottom row represents the results from simulations with OBC implicit model pair with the same force field.

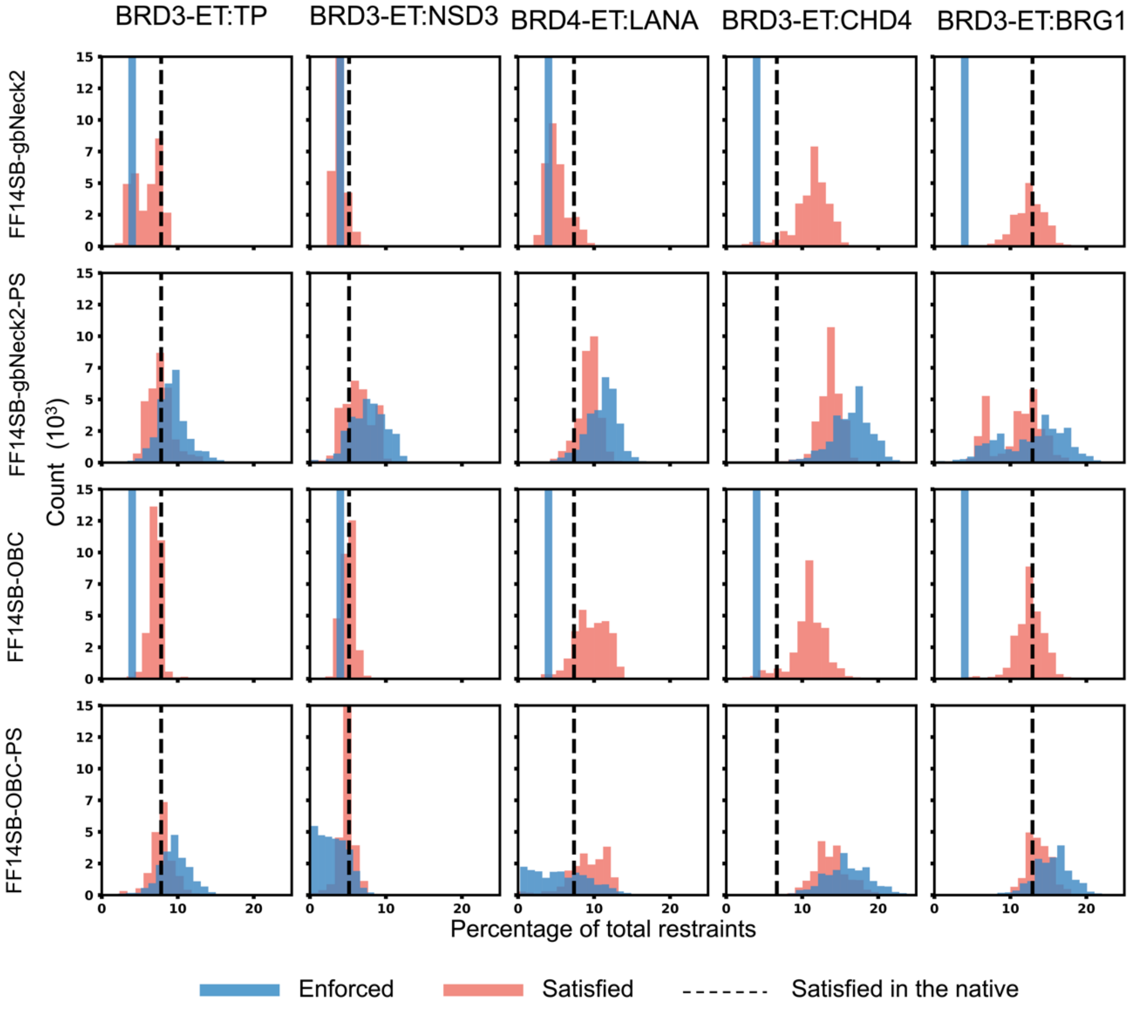


**Figure S17. Restraint distributions for different protein-peptide complexes under a series of force field.** For all 5 test cases, the distribution of enforced and satisfied contacts in the lowest temperature replica is plotted here along with the number of satisfied contacts in the experimental structure. The top row represents results from the simulations with FF14SBside force field with gbNeck2 implicit model with the vanilla MELD approach, whereas the second row represents the results from simulations with MELD-Adapt approach (with parameter sampling) under similar physical model as the top row. Similarly, the third row represents results from the simulations with FF14SBside force field with OBC implicit model with the vanilla MELD approach, whereas the fourth row represents the results from simulations with MELD-Adapt approach (with parameter sampling) under similar physical model as the third row. All MELD-Adapt simulations were run with a fixed reward value of 0.25 kcal/mol.

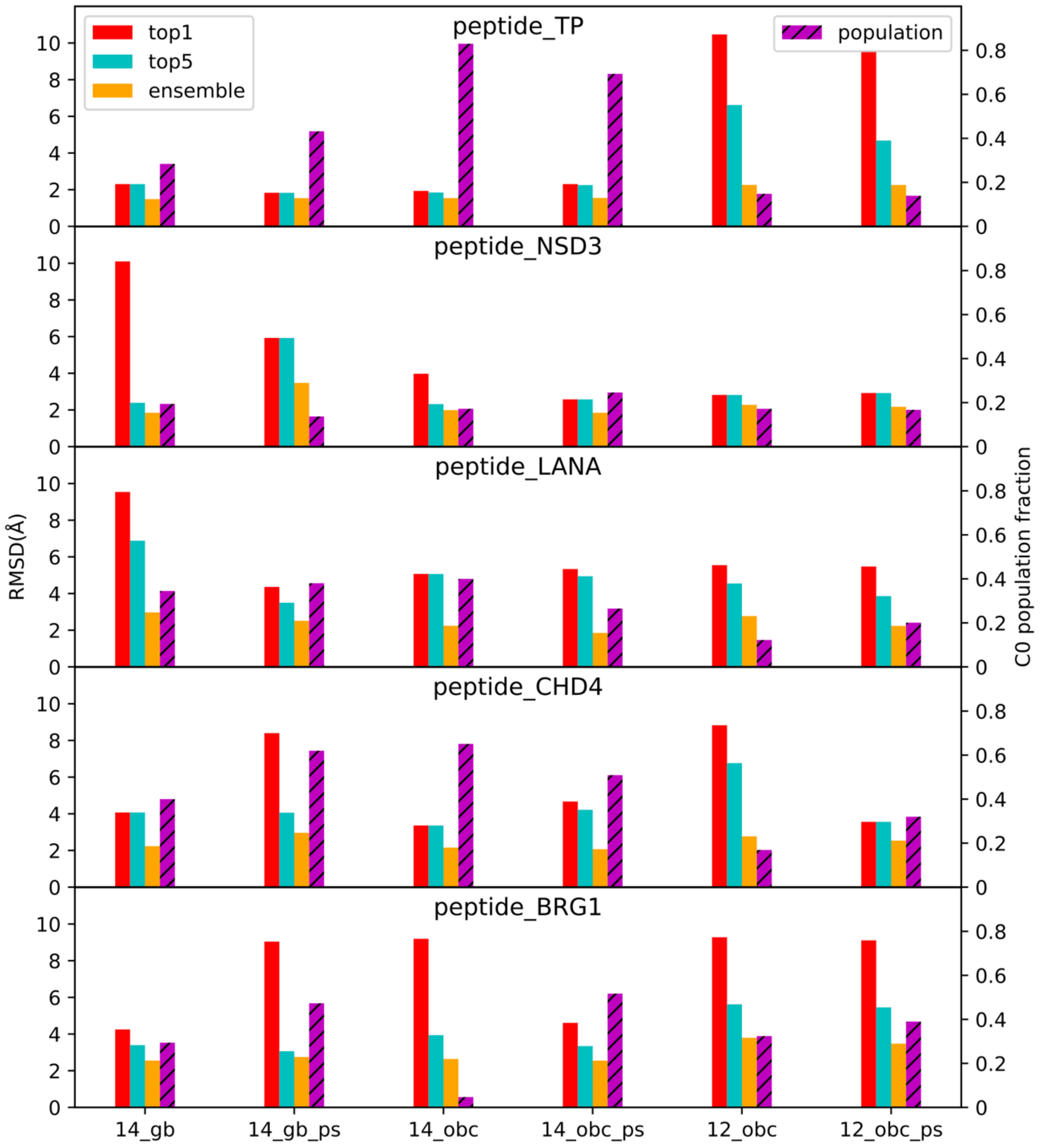


**Figure S18. Quantitative analysis of the benchmark set under different condition.** For all 5 test cases, the RMSD of top1 cluster centroid, best of top5 cluster centroid and best the lowest temperature replica with respect to the experimental structures is plotted here along with the population of the top clusters. X axis represents 6 different force field condition that we tested: FF14SBside force field with gbNeck2 implicit model with the vanilla MELD approach, MELD-Adapt approach (with parameter sampling) under similar physical model as the previous approach, FF14SBside force field with OBC implicit model with the vanilla MELD approach, MELD-Adapt approach (with parameter sampling) under similar physical model as in the previous one. All MELD-Adapt simulations were run with a fixed reward value of 0.25 kcal/mol.
